# Sampling antibody conformational ensembles with ABodyBuilder4-STEROIDS

**DOI:** 10.64898/2026.04.14.718378

**Authors:** Fabian C. Spoendlin, Matteo Cagiada, King Ifashe, Odysseas Vavourakis, Charlotte M. Deane

## Abstract

Conformational flexibility is fundamental to the function of many proteins and in the case of antibodies can impact key properties such as affinity and specificity. While it is possible to predict single, static protein structures with high accuracy, predicting conformational ensemble remains challenging. Molecular dynamics simulations suffer from high computational costs, while deep learning methods are yet to achieve the same level of accuracy. Here, we introduce ABB4-STEROIDS a generative structure prediction model that samples conformational ensembles of antibodies. We trained our model on 4.2 million structural frames derived from ∼136,000 coarse-grained and a set of 83 new all-atom antibody MD simulations. We benchmarked our model on reproducing MD ensembles and evaluated the diversity of sampled structures and the covered conformational space against experimental evidence. ABB4-STEROIDS achieves state-of-the-art accuracy, particularly within the experimental benchmarks. The model is openly available and provides a robust resource for large-scale investigations of antibody conformational ensembles.

## 1 Introduction

Conformational changes are fundamental to the function of many proteins enabling dynamic processes such as binding, allostery and signalling events [1, 2]. In antibodies specifically, the majority of structural diversity is concentrated in the complementarity determining regions (CDRs) [3, 4], while the frameworks remain more static [5]. CDR flexibility has been linked to key functional properties such as binding specificity and affinity. First, structural diversity can allow a single antibody to bind a range of antigens in distinct conformational states [6, 7, 8]. Second, flexibility affects the thermodynamics of binding influencing both enthalpic and entropic contributions [9]. Several studies on individual antibodies have found that antibody maturation is associated with CDR rigidification [10, 11, 12, 6, 13, 14] which suggest that modulating flexibility is a natural mechanism of the immune system to balance between more promiscuous naive and higher affinity mature antibodies. However, larger scale studies have not seen a similar trend at the repertoire level [15, 16].

As affinity and specificity are critical for antibody drug development, characterising the conformational landscapes of CDRs would provide valuable insights [17]. To this end, methods to classify whether CDRs are able to undergo conformational changes are available [18]. However, modelling exact structures of conformational ensembles remains challenging.

Molecular dynamics (MD) simulations are a powerful tool to study conformational changes. However, standard all-atom MD is constrained by high computational costs. Even with specialised supercomputers [19] or enhanced sampling [20] it remains challenging to simulate large number of proteins over long timescales. Therefore, large-scale datasets of all-atom MD simulations are limited to approximately thousands of proteins simulated for hundreds of nanoseconds, restricting both the exploration of protein space and coverage of conformational landscapes [21, 22]. Coarse-grained simulations provide a more efficient alternative but they can suffer from reduced accuracy introduced by the approximation of the underlying force field [23, 24]. One-bead-per-residue models, such as CALVADOS [25], have been used to create datasets as large as tens to hundreds of thousands of proteins [26, 16]. For example, the Flexibility Antibody Database (FlAbDab) [16] contains simulations of more than 150,000 antibody Fv domains.

Machine learning methods offer the potential of even more efficient sampling of conformational ensembles. Initial uses of ML in the flexibility space focussed on stochastic subsampling of the multiple sequence alignment (MSA) used by AlphaFold2[27] during inference to increase output diversity [28, 29, 30, 31, 32, 33, 34, 35]. Recent structure predictors employ generative models which are in theory better suited for sampling distributions of conformational states. However, models such as AlphaFold3 [36] and Boltz-1 [37] are trained on crystal structure data only and are limited in their ability to sample diverse conformations [36, 37]. To address this, a range of specialised models trained on MD data has emerged which demonstrate improved modelling of conformational ensembles [38, 39, 40, 41]. AlphaFlow [40] and BioEmu [38] are regular structure predictors that map an input sequence to an ensemble of structures, whereas aSAM [39] takes both initial structure and sequence as input to produce alternative conformations.

Guidance and steering of structure predictors at inference time towards lower-resolution experimental constraints has shown promising results for sampling desired conformational states. For example, the ROCKET method [42] optimises MSA profiles to fit output structures to cryo-EM maps of a target state. Similarly, the ConforMix method uses a constraint-based guidance function during the reverse diffusion process to sample a conditional distribution and steer predictions towards a set of constraints associated with the target state [43].

While ML-based conformation predictors have shown promising results in modelling general proteins, no method trained and evaluated on antibody data alone is available. To address this gap, we present **A**ntibody**B**uilder**4** - **S**tructure predictor **T**uned on **E**nsembles of complementary determining **R**egions **O**bserved **I**n molecular **D**ynamics **S**imulations (ABB4-STEROIDS) a generative structure prediction model that samples conformational states of antibodies (Figure 1a). ABB4-STEROIDS is a flow matching model pre-trained on 4.2 million representative frames of ∼ 136, 000 coarse-grained antibody MD simulations from the FlAbDab [16] and fine-tuned on a new set of 83 all-atom MD simulations, which we release alongside the model (https://doi.org/10.5281/zenodo.19471889). We demonstrate state-of-the-art performance at reproducing ensemble metrics observed in the MD simulations and matching the flexibility and conformational space of antibodies observed in experimental data. ABB4-STEROIDS is openly available on Github (github.com/oxpig/ABB4).

**Figure 1:**
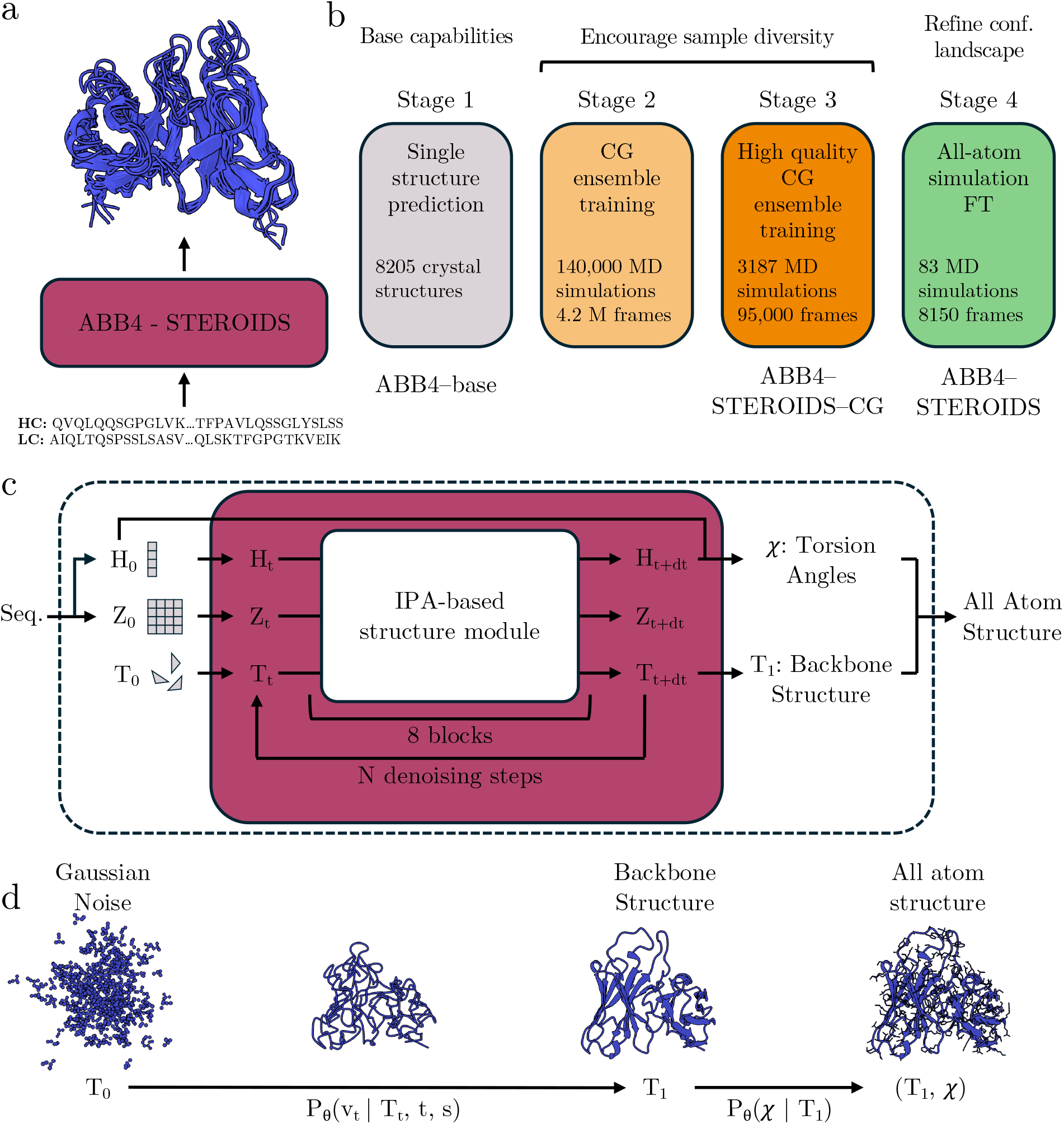
Overview of the ABB4 models. a) ABB4-STEROIDS generates an ensemble of antibody structures for a given input sequence. b) Summary of the four-stage training procedure. c) Illustration of the model architecture. *H*: single/node representation encoding residue level features. *Z*: pair/edge representation encoding inter-residue features. d) Diagram providing an overview of the flow matching methodology. *T* : backbone frames *χ*: Torsion angles. Subscripts denote the flow matching time step.

## 2 Results

### 2.1 Overview of the ABodyBuilder4 models

ABB4-STEROIDS is a generative structure prediction model trained to sample conformational ensembles of antibodies (Figure 1a). In this section, an overview of the model architecture and training procedure is presented (for details see methods and SI).

The architecture builds on the AF2 [27] structure module (Figure 1c). An input sequence is embedded into a single representation (*H*) and a pair representation (*Z*). Protein structures are represented by a frame parametrisation (*T*) where all backbone atoms of a residue are modelled by a single rotation matrix and translation vector. The three representations are updated by 8 blocks of a structure module using Invariant Point Attention (IPA) at its core and side chain torsion angles (*χ*) are predicted with an additional head (Figure 7).

ABB4 is trained as a flow matching model over backbone frames largely following the Frame-Flow [44] methodology (Figure 1d). During training, a target structure and a time step are sampled and the corresponding noise applied to the backbone frames to obtained a partially noised structure (*T*_*t*_). Conditioned on noised backbone frames (*T*_*t*_) the model is trained to predict the clean backbone (*T*_1_) and torsion angles (*χ*). During inference, initially pure noise is sampled (*T*_0_). From model predictions a vector field (v_t_) is constructed that interpolates between *T*_*t*_ and *T*_*t*+*dt*_ and denoises the samples. After a full model rollout a clean backbone structure is reconstructed from the final frame (*T*_1_). Torsion angles (*χ*) predicted during the final inference step are used to build an all-atom structure.

We implemented a novel four-stage training strategy to optimise conformation sampling (Figure 1b). In stage 1, a model was trained on experimental structures to establish basic single structure prediction capabilities. We refer to the resulting model as ABB4-base which serves as a foundation for subsequent stages. In stages 2 and 3, the base model was pre-trained on a large corpus of coarse-grained MD simulation data deposited in the FlAbDab [16]. The objective of these stages was to encourage the model to sample diverse conformational ensembles, although the model may inherit biases of the coarse-grained force field. More specifically, stage 2 utilised the complete dataset, comprising over 4.2 million structural frames extracted from ∼ 136, 000 antibody simulations, including both high-quality trajectories initiated from experimental structures and lower-quality simulations initiated from predicted structural models. In stage 3, the training was restricted exclusively to a high quality subset of FlAbDab. The resulting model from stages 1 to 3 is designated ABB4-STEROIDS-CG. In stage 4, we created our final ABB4-STEROIDS model by fine-tuning ABB4-STEROIDS-CG on the smaller set of 83 all-atom simulations. The primary objective of this stage was to mitigate biases introduced through the coarse-grained data and to refine the sampled landscape and relative populations of conformational states.

### 2.2 ABodyBuilder4-base serves as a strong base for structure prediction

We benchmark the ABB4-base model for single structure prediction against open-source antibody specific and general protein structure prediction models. We compared against ABody-Builder3 (ABB3) [45], an antibody specific model based on the AF2 architecture and a regression framework which was trained on isolated antibody structures (agnostic of the antigen). We also evaluated Boltz-1 [37] as a representative of AF3-style generative models trained on the full PDB including protein complexes.

We compared the performance of the three methods on a reduced test set of 91 antibodies (Figure 2). Our complete test set of 100 antibodies was curated to exclude any antibody with more than 80% sequence identity (calculated across concatenated CDRs) to structures released prior the training cut-off of the benchmarked methods. An exception is ABB3, which did not use a cut-off-based data split, and consequently, to prevent data leakage, we removed all entries overlapping with the ABB3 training set from out test set when evaluating this specific model. A comparison of ABB4-base and Boltz-1 on the full test set is provided in Figure S5.

**Figure 2:**
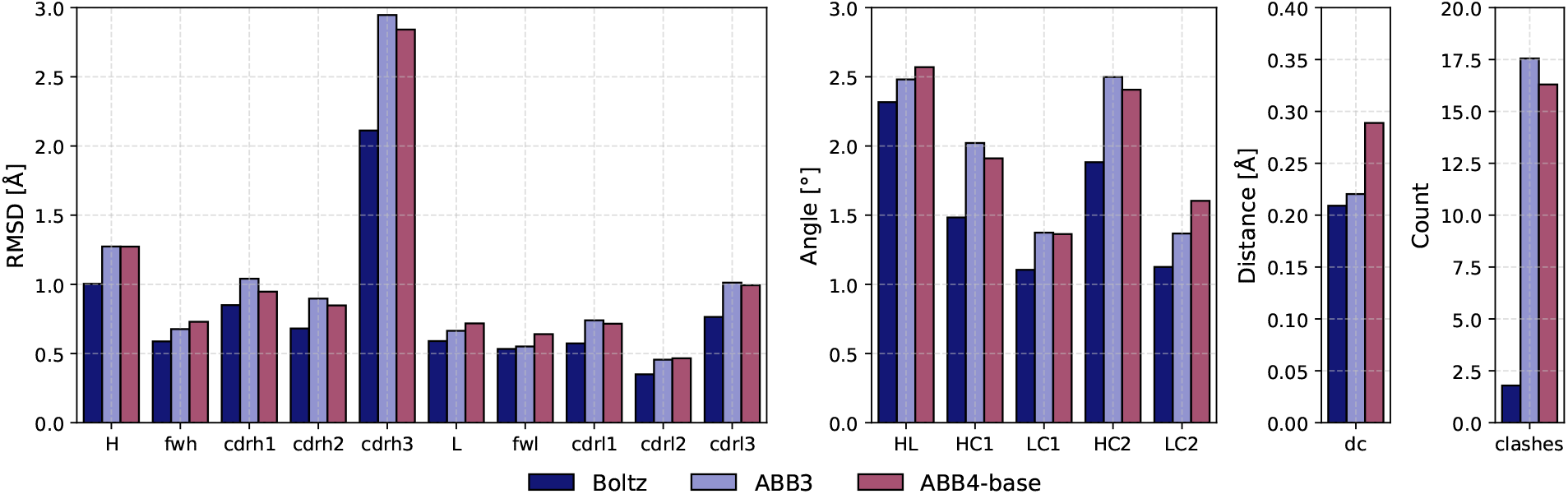
Evaluation of ABB4-base single structure prediction against Boltz-1 and ABB3 on a reduced test set of 91 antibody structures. Predictions were assessed using RMSD to the crystal structure of all antibody regions individually (H: heavy chain, fwh: heavy chain framework, L: light chain, fwl: light chain frame work), deviations in VH-VL orientation angels (HL, HC1, LC1, HC2, LC2) and distance (dc) as defined in [46] compared to the crystal structure and the number of atomic clashes defined as an overlap in van der Waals radii of any backbone or side chain atoms.

Structure prediction accuracy was assessed using the RMSD and differences in VH-VL orientations compared to the crystal structure as well as physical validity measured by atomic clashes. Across all metrics, we observed a general trend of ABB4-base showing small improvements over ABB3 and Boltz-1 providing a more substantial advancement. These results highlight that while ABB4-base does not achieve state-of-the-art accuracy at single structure prediction, it can serve as a reliable base model which is competitive with contemporary structure predictors.

### 2.3 ABodyBuilder4-STEROIDS accurately samples antibody conformational ensembles

We next trained the ABB4-base model on MD data to obtain ABB4-STEROIDS-CG and ABB4-STEROIDS. We evaluated the models on held-out MD test sets and compared against a set of experimentally observed ensembles.

We benchmarked our models against established structure and ensemble prediction methods. From the former group we included Boltz-1 [37] and AF2 with MSA subsampling, a workflow generating higher-diversity samples (e.g. [30]). From the latter group, we used aSAM [39], AlphaFlow [40] and BioEmu [38]. We generated 100 structures for each antibody with all of the assessed models to evaluate ensemble prediction.

#### 2.3.1 Predictions accurately reproduce MD ensemble metrics

In a first step, we compared model predictions against the coarse-grained MD data on the full test set of 100 antibodies and the all-atom MD test set of 14 antibodies. We focused our analysis on ensemble diversity metrics (ensemble RMSD & RMSF) to evaluate the magnitudes of flexibility in the predicted ensembles. We further report metrics based on the distances between predicted and MD frames (recall, precision and distributional accuracy).

The ensembles generated by the seven methods show large differences in the spread of the sampled structures, reflecting the predicted flexibility (Figures 3a, 4a&b, S7a, S8a&b). Collectively across all CDRs, our ABB4-STEROIDS-CG model generates the most diverse ensembles whereas our ABB4-STEROIDS model predicts slightly reduced flexibility in CDR1s and CDR3s, with a more pronounced reduction observed in CDR2s. All other methods generally underestimate the flexibility of canonical CDRs. In particular, Boltz-1 ensembles display very low ensemble RMSD and RMSF values. Across all models, higher diversity is predicted for CDRH3s. Dissociation of the VH and VL domains is rare, even for methods not explicitly trained on multi-domain proteins (such as AlphaFlow and aSAM), with the exception of BioEmu, where dissociation occurs more frequently (Figure S6).

**Figure 3:**
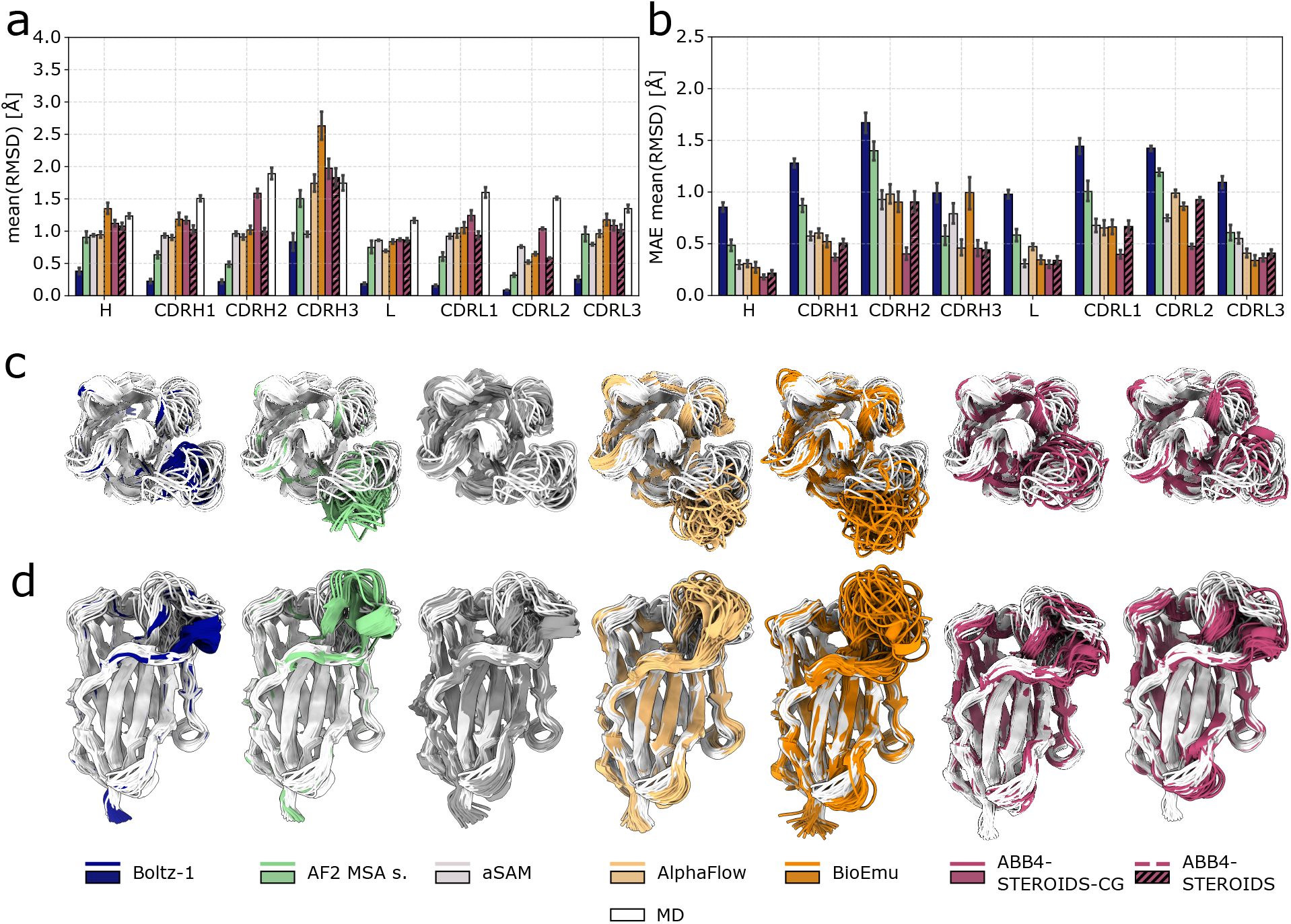
Evaluation of predicted structural ensembles against coarse-grained MD data. a) Mean ensemble RMSD in predictions and MD trajectories across all antibodies in the coarse-grained MD test set. Values computed across the heavy chain (H), light chain (L) and individual CDRs. b) Mean absolute error (MAE) of ensemble RMSD in the predictions compared to the MD trajectories. Top (c) and side (d) view of predicted ensembles for antibody PDB 9DS1 overlaid with coarse-grained MD frames. Only antibody heavy chains are shown, light chains were removed for visualisation.

**Figure 4:**
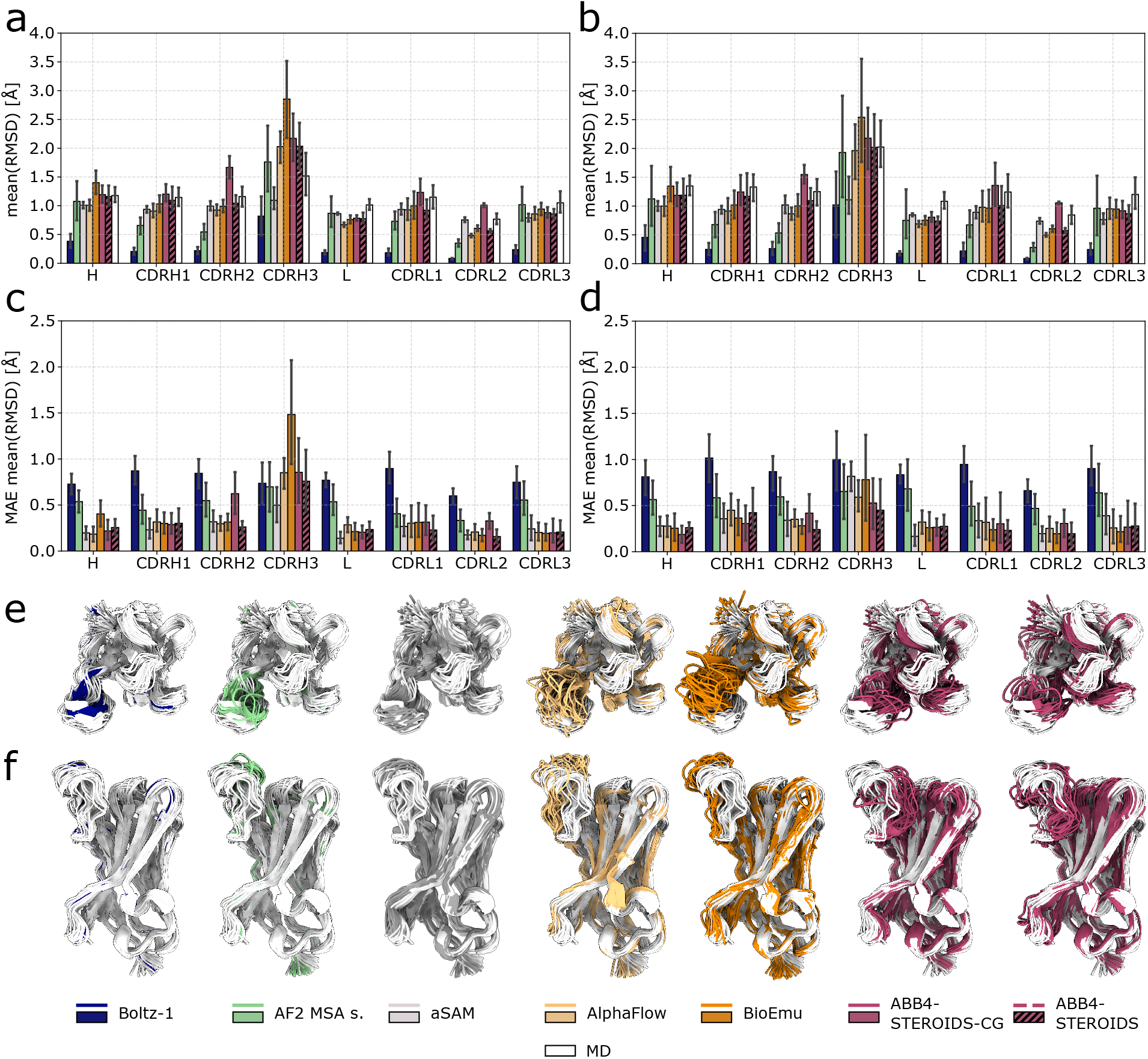
Evaluation of predicted structural ensembles against all-atom MD data. Mean ensemble RMSD in predictions and MD trajectories (a) across all antibodies in the all-atom test set (N=14) and (b) across antibodies with flexible CDRH3s (ensemble RMSD > 1.4 Å in MD trajectories; N=7). Values computed across the heavy chain (H), light chain (L) and individual CDRs. Mean absolute error (MAE) of ensemble RMSD in the predictions compared to the all-atom MD data for (c) all antibodies and (d) antibodies with flexible CDRH3s. Top (e) and side (f) view of predicted ensembles for antibody PDB 8VVB overlaid with all-atom MD frames. Only antibody heavy chains are shown, light chains were removed for visualisation.

We quantified the agreement between predicted flexibility and MD by computing the errors in ensemble RMSD and RMSF. Within the coarse-grained test set (Figure 3b and Figure S7b), Boltz-1 and AF2 MSA subsampling exhibit the highest errors overall, while aSAM, AlphaFlow and BioEmu tend to perform better. ABB4-STEROIDS-CG consistently shows the highest accuracy in both metrics across all antibody domains and ABB4-STEROIDS maintains comparable performance for CDR1s and CDR3s, though it shows larger errors for CDR2. Increased CDR2 errors in the later case may originate from a known limitation of the coarse-grained MD force field, which tends to overestimate CDR2 flexibility [16].

On the all-atom test set, ABB4-STEROIDS achieved top or near top performance for all canonical CDRs while Boltz-1, MSA subsampling, and aSAM perform well on CDRH3 when measured across the full set (Figure 4c and Figure S8c). Our analysis reveals that these models, particularly Boltz-1 and aSAM, predict ensembles with low diversity that is largely independent of the true CDRH3 flexibility (Figure S8f). As the all-atom test set contains several antibodies with relatively static CDRH3s these methods attain high scores. We further evaluated performance on a refined subset excluding antibodies with low CDRH3 flexibility (ensemble RMSD > 1.4 Å). On this more challenging set, the accuracy of Boltz-1, MSA subsampling, and aSAM dropped drastically, whereas ABB4-STEROIDS emerged as the most accurate predictor for CDRH3s (Figure 4d and Figure S8d). These observations suggest that while several methods are competitive at modelling antibodies with static CDRs, ABB4-STEROIDS is most capable of capturing the dynamics of flexible antibodies.

For completeness, we computed metrics of recall, precision and distributional accuracy (see Methods for a definition of metrics). However, we note important weaknesses of these metrics to evaluate ensemble prediction. Recall measures how closely each MD conformational state is covered but does not penalise excessive conformational entropy. Furthermore, the metric is agnostic to the temporal dynamics of the MD trajectories and therefore high-frequency structural fluctuations are weighted equally to the slower transitions that typically characterize functional states. Precision is defined as the minimum distance from each structure of the predicted ensemble to any MD frame and does not provide any information on how accurately an ensemble is reproduced. Distributional accuracy indicates the Wasserstein distance between the predicted and MD distributions.

ABB4-STEROIDS and ABB4-STEROIDS-CG show good recall values for canonical CDRs (Figure S7c and Figure S8e). For CDRH3, all methods exhibit lower recall and several outperform the ABB4-STEROIDS models. Metrics of precision and distributional accuracy are dominated by Boltz-1 and AF2 MSA which are highly accurate structure predictors but generate ensembles of low flexibility which do not match the simulations (Figure S9).

Overall, the ABB4-STEROIDS models accurately reproduce the diversity of structural ensemble observed in MD simulations, as evidenced by ensemble RMSD and RMSF metrics. As expected, ABB4-STEROIDS-CG demonstrates superior performance on the coarse-grained test set, while the ABB4-STEROIDS model exhibits higher accuracy on the all-atom test set. The large size of the coarse-grained set provides a more robust estimate of performance, albeit with the caveat that the underlying conformational distributions may inherit known biases, particularly for CDRH2 [16]. In contrast, the all-atom set is more reliable but necessitates a cautious interpretation of results due to the limited size.

#### 2.3.2 All-atom MD fine-tuning improves conformational sampling and physical validity

The coarse-grained MD data used in pre-training has inherent limitations. First, the CALVA-DOS 3-Fv parametrisation that was used to generate these data was optimised to reproduce all-atom ensemble averages. Consequently, certain conformational states may be under-sampled or their conformational distributions may differ from those of more accurate all-atom simulations [16, 23, 24]. Second, the force field models each residue as one bead. Coordinates of individual atoms are reconstructed with an additional model which may introduce interatomic clashes [16]. A short fine-tuning stage on a small set of all-atom simulations (training stage 4) was introduced to prevent our final ABB4-STEROIDS model from inheriting these biases. This fine-tuning improves the sampled conformations and enhances the physical validity of generations.

We performed time-independent component analysis (tICA) to compare conformational landscapes with the all-atom MD simulations. tICA reports on the slowest structural transitions in the simulations [47] which are typically the most biologically relevant overcoming some of the limitations we previously discussed for the recall metric.

Fine-tuning on all-atom simulations enables ABB4-STEROIDS to capture the conformational space of antibodies more accurately than ABB4-STEROIDS-CG. Several case studies of test set antibodies revealed two modes of refinement (Figure 5a). First, for CDRs poorly predicted by ABB4-STEROIDS-CG a larger fractions of predictions shifts into MD-occupied regions (e.g. CDRH3 of PDB 8r1d in Figure 5a). Second, for CDRs where ABB4-STEROIDS-CG primarily samples within the MD distributions, we frequently observed more accurate weighting of equilibrium densities. Specifically, the fine-tuned model reduces the tendency of the coarse-grained model to over-represent rare conformational states (e.g. CDRH3 of PDB 8w83 and CDRH2/CDRL1 of PDB 8f2t in Figure 5a).

**Figure 5:**
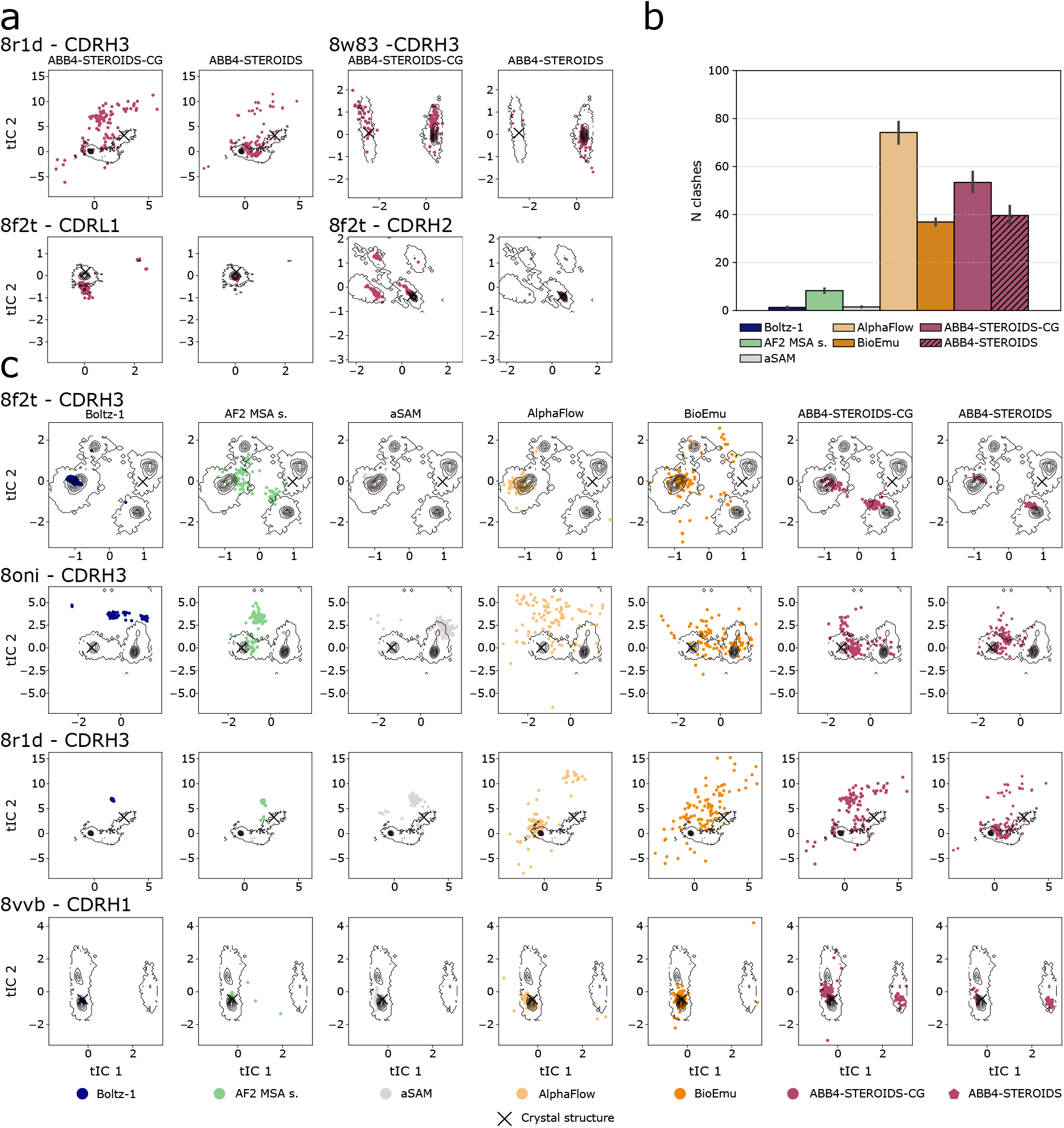
Sampling of conformational states and physical validity of predictions. a) Comparison of the conformational ensemble sampled by ABB4-STEROIDS-CG and ABB4-STEROIDS using tICA. b) Average number of atomic clashes in predicted structural model of the test set. c) Comparison of the conformational ensemble sampled by all seven benchmarked methods. a & c) The first two time-lagged independent components (tICs) are plotted. The black contour indicates the all-atom MD distributions. Predicted structures were mapped to the tICs and are represented by the coloured dots. The crystal structure from which the simulations was started is shown by the black cross.

Our comparison of all seven benchmarked methods suggests that ABB4-STEROIDS predictions most closely match the all-atom MD distributions in tICA space (case studies shown in Figure 5c; full test set in Figures S10-S22). This is exemplified by the CDRH3 of PDB 8f2t (Figure 5c), where ABB4-STEROIDS is the only model to effectively capture two major equilibrium states. The other methods either tend to sample a single state or generate large number of structures outside the MD distributions. Similarly, for CDRH3 of PDB 8oni and 8r1d (Figure 5c), where most benchmarked methods predict ensembles largely disjoint from the ground truth, ABB4-STEROIDS recovers the largest fraction of the MD conformational space. Finally, for CDRH1 of PDB 8vvb (Figure 5c), ABB4-STEROIDS uniquely captures a second conformational state omitted by all other methods.

We assessed the physical validity of model predictions by quantifying inter-atomic clashes (Figure 5b). Comparison between the ABB4 models show that the all-atom fine-tuning reduces steric violations. In the broader benchmark, ABB4-STEROIDS exhibits a clash profile comparable to BioEmu, whereas AlphaFlow displays the higher frequencies. As expected, static predictors (Boltz-1 and AF2 MSA subsampling) produce the fewest violations. Among the ensemble-based methods, only aSAM achieves similar physical validity. However, this performance is likely influenced by its reliance on a high-quality Boltz-1 input template.

#### 2.3.3 Ensembles are consistent with experimental evidence on flexibility

After extensive evaluation against MD ensembles, we compared our predictions with experimental data. To this end, we collected and compiled ensembles of antibodies derived from multiple experimental structures. However, such data has limitations as experimental structures do not necessarily resolve all physiologically relevant conformational states and do not inform on temporal trajectories. We, therefore, focussed our comparison on assessing the overall predicted flexibility and coverage of the conformational states found in experimental ensembles using a set of metrics we introduced in previous works [18, 16]. Furthermore, because we could not assemble a sufficiently large dataset consisting exclusively of structures released after the training cut-offs, the test set used in this section includes antibodies that overlap with the training sets of all the methods (see Methods for details).

We assessed the correspondence between predicted ensemble diversity and experimental flexibility. Given that calculating average RMSD values for incomplete experimental ensembles is unreliable, we adopted a binary classification approach. We collected all antibodies in SAbDab represented in multiple structures [48, 49] and categorised them as ‘rigid’ and ‘flexible’ based on whether structural evidence supported a single or multiple accessible conformations (see Methods for details). We then quantified the agreement between the RMSD in predicted ensembles and these binary labels using a point-biserial correlation (Figure 6a). The ABB4 models achieved the highest correlation across all CDRs among the benchmarked methods. Similar to previously evaluated metrics, we observed an improved correlation for CDRH2 of ABB4-STEROIDS over ABB4-STEROIDS-CG.

**Figure 6:**
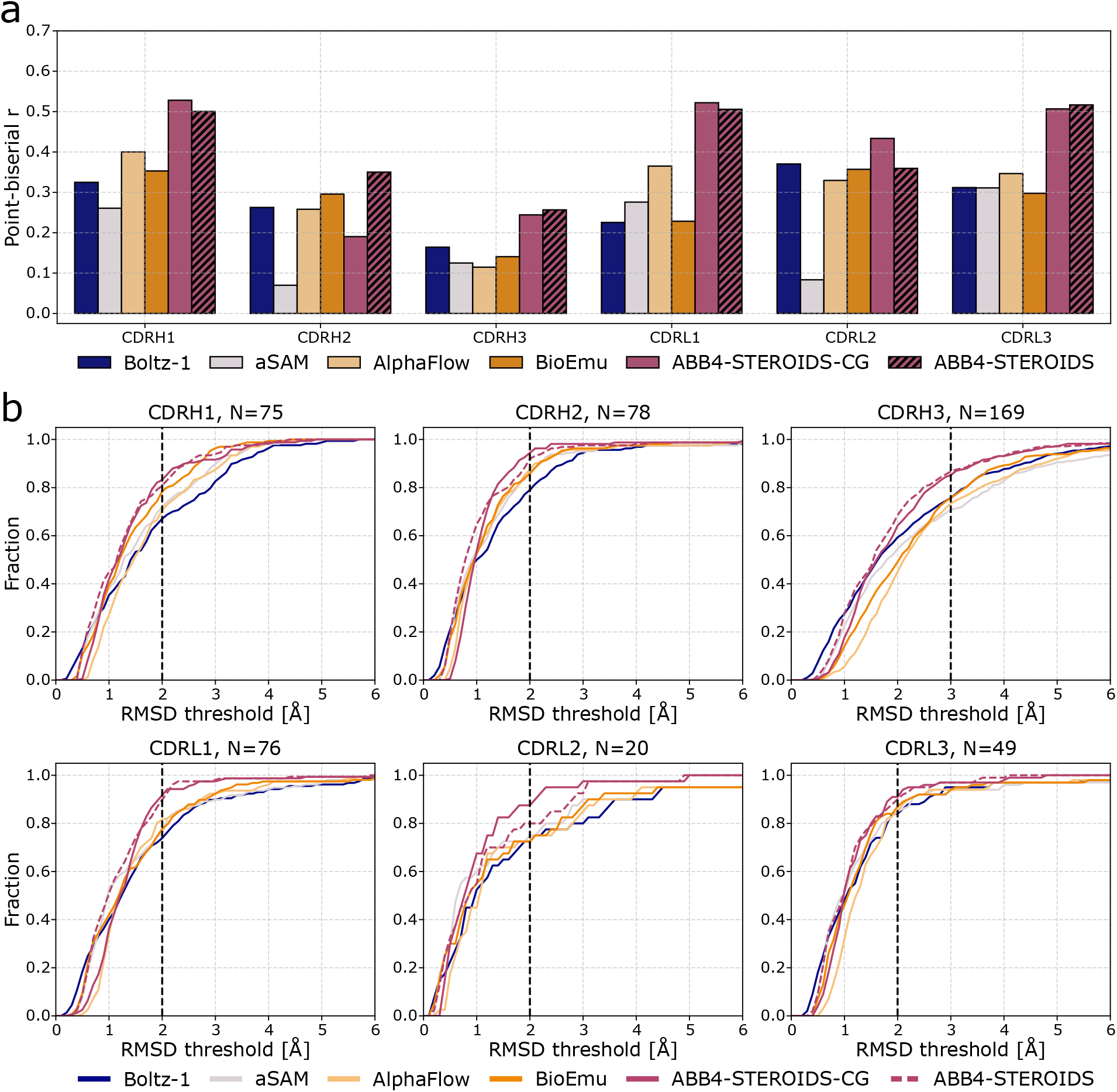
Consistency of predictions with experimental evidence of flexibility. a) Flexibility correlation. Point-biserial correlation coefficient (r) between binary flexibility labels assigned to experimental ensemble and RMSD in predictions for all CDR loops. b) Conformation coverage. Fraction of experimental conformations covered by model predictions as a function of distance. Dashed lines highlight ABB4-STEROIDS/-CG covering the highest number of conformations at given distance thresholds.

We further evaluated the coverage of experimental conformations by the predictions. To compute this metric, we gathered all antibodies labelled as ‘flexible’ (multiple conformational states) in the previous section and calculated how closely each experimental conformation is covered by a prediction (Figure 6b). Boltz-1 covers a small fraction of conformations with very low RMSD (even below the experimental error margins) showing that it efficiently reproduces some of the crystal structures. However, its limited diversity prevents capturing of the broader ensemble. aSAM, AlphaFlow, BioEmu and ABB4 models outperform Boltz-1 at covering larger fractions of conformations. Among these, the ABB4 models consistently define the Pareto front, demonstrating the best balance between structural accuracy and diversity. More specifically, ABB4 models cover the highest number of conformations for canonical loops at 2 Å and for CDRH3 at 3 Å.

#### 2.3.4 Ablation studies and model variants

Following the evaluation of the main models, we assessed several model ablations and variants. To ensure results are robust, we conducted these evaluations on the larger coarse-grained MD rather than the small all-atom test set. All performance metrics are benchmarked against the ABB4-STEROIDS-CG model as the primary baseline.

As the number of inference steps directly scales computational cost, we evaluated the relation ship between sampling depth and model performance (Figure S24). Reducing the default 100 steps to 50 has a negligible impact on performance. A slight decrease in accuracy is observed at 20 and 10 steps, followed by a loss of performance at 5 steps. These results suggest that the inference steps can be halved to 50 steps without compromising performance, though further reductions incur a progressively higher cost.

We further examined the impact of training stage 2, which incorporates a large volume of potentially lower quality coarse-grained MD simulations initialised from predicted structural models. An ablated model variant trained excursively with stages 1 and 3 (ABB4-STEROIDS-1,3; Figure S23) performs slightly better on MD-based validations. The lack of difference between ABB4-STEROIDS-1,3 and ABB4-STEROIDS-CG despite the larger training data of the later likely stems from subtle differences in CDR dynamics between the simulations initialised from models versus experimental structures (SI Section S2.2). However, including stage 2 leads to clear performance gains on the experimental benchmark (Figure S23). This suggests that the larger, more diverse dataset prevents the model from overfitting on the MD distributions and enhance generalisation to experimental ensembles.

Finally, we explored a model variant incorporating an additional loss function targeted at ensemble properties. Models for conformational sampling are typically trained using standard structure prediction losses applied to individual frames. While training frames are sampled from MD trajectories to capture dynamics, the objective is computed on a single prediction at a time. To better align the training objective with ensemble prediction, we introduced an ensemble-based loss function that to minimises the error in residue-level RMSF (SI Section S1.1.2). We observed some training instability when the difference between predicted and ground-truth RMSF was high, though this was partially mitigated by employing a *χ*^2^-style loss instead of a standard Mean Squared Error (MSE). Inclusion of this additional loss improved ensemble metrics for canonical CDRs (Figure S25). However, performance for CDRH3 was slightly reduced, likely due to the inherent difficulty in predicting the RMSF of this loop which further increased the training instabilities associated with this loss. While the ensemble loss is effective for domains where RMSF can be predicted with reasonable accuracy, further refinement is needed for regions with higher uncertainties.

## 3 Discussion

While predicting a single, static structure is now possible with high accuracy for the majority of globular and monomeric proteins, accurately capturing the conformational heterogeneity of these molecules remains a significant challenge. Several approaches have been explored to predict conformational ensembles. One set of methods focussed on inference time modification of common structure predictors such as MSA subsampling, guidance and steering [30, 42, 43]. Alternative approaches trained specialised ensemble prediction models using MD data [39, 40, 38]. Here, we introduced ABB4-STEROIDS a generative structure prediction tool that samples conformational ensembles of antibodies with high accuracy.

In a first step, we trained ABB4-base as a standard antibody structure predictor. Following the antigen-agnostic paradigm of previous ABodyBuilder iterations (ABB2, ABB3), which unlike ABB4 are based on AF2-style regression models [45, 50], we trained exclusively on antibody structures (removing antigens when present). This contrasts with recent AF3-style generative models like Boltz-1 [37] that incorporate antibody-antigen complexes.

The generative framework of ABB4-base resulted in small improvements over ABB3 [45] and Boltz-1 achieves even higher overall accuracy. While it is well established that AF3-style models are better for modelling antibody-antigen complexes [36, 51], our results substantiate more tentative evidence suggesting that they may also be more accurate than AF2-style alternatives for antigen-agnostic structure prediction [52, 53]. The marginal performance gains observed across several generations of antigen-agnostic model (ABB2, ABB3, ABB4-base, AF2) [27, 50, 45], suggest a possible accuracy ceiling for this training regime. We, therefore, speculate that protein complexes may be informative for predicting antibody structures, especially given that most antibodies in the PDB are resolved bound to their antigen.

ABB4-base serves as an accurate base model which we trained further on a large dataset of coarse-grained antibody MD simulations to obtain ABB4-STEROIDS-CG. For these training stages, we used simulations deposited in the FlAbDab [16] from which we extracted over 4.2 million representative trajectory frames capturing ∼ 136, 000 antibodies. In a final stage, the model was fine-tuned on a new set of 83 antibody all-atom simulations, which we release alongside the model, to obtain ABB4-STEROIDS.

We evaluated the magnitudes of flexibility in the predicted ensembles against coarse-grained and all-atom MD data. The larger scale of the coarse-grained set (N=100) gives a more robust estimate of performance. However, the underlying conformational distributions may inherit force field biases. Dynamics of the the all-atom set are more reliable but the limited size (N=14) requires cautious interpretation of results. Overall, ABB4-STEROIDS-CG is slightly more accurate on the coarse-grained test set, while the ABB4-STEROIDS model exhibits marginally lower errors on the all-atom test set. A significant discrepancy in performance was observed only for the CDR2 loops. This is likely attributed to a known limitation of the coarse-grained modelling protocol (CALVADOS 3-Fv) overestimating the flexibility of these CDRs [16]. This results also suggests that all-atom MD fine-tuning mitigates biases in CDRH2 modelling introduced through the coarse-grained data [16].

Our ABB4-STEROIDS/-CG models outperform all alternative methods we assessed (Boltz-1 [37], AF2 MSA subsampling [28, 30, 29], aSAM [39], AlphaFlow [40], BioEmu [38]) when comparing magnitudes of flexibility against both types of MD simulations. The performance gains are more pronounced for canonical CDRs for which most methods tend to substantially underestimate structural diversity. Improvements for CDRH3s are smaller, which we link to the magnitude of CDRH3 flexibility being on a similar scale to current prediction accuracy [53, 45]. This inherent uncertainty increases output diversity across all models and makes accurate prediction of conformational states particularly challenging. Nevertheless, when evaluated against the coarse-grained test set, our models achieved top performance. Analysis against the all-atom set revealed that while methods such as Boltz-1 and aSAM remain competitive at modelling static loops, ABB4-STEROIDS provides a superior representation of more dynamic CDRH3s.

Other than CDRH2 flexibility, the coarse-grained MD data exhibits two further inherent limitations. First, coarse-grained models are known to smooth the protein energy landscape [23, 24], which can lead to an imprecise modelling of conformational distributions. Second, individual atomic coordinates need to be reconstructed which can result in interatomic clashes [16]. The introduction of a short fine-tuning stage on a small number of all-atom simulations mitigated both of these issues and ABB4-STEROIDS demonstrated improved sampling of conformational distributions and fewer steric violations than ABB4-STEROIDS-CG.

Comparing against alternative methods showed that ABB4-STEROIDS most accurately samples the conformational landscape observed in all-atom MD simulations. However, we also show that specific conformational states can be missed and relative population densities may not be accurate. These findings underpin that sampling conformational ensembles with structure prediction model remains a challenging task.

As MD may have its own inaccuracies and biases we compared predictions against experimental data which serves as our most critical benchmark. Following a previously established protocol [18], we curated a dataset from SAbDab [48, 49] consisting of antibodies with multiple resolved structures. Initially, we categorised these antibodies using binary labels to distinguish between those with single versus multiple accessible conformations. Correlating these labels with the predicted ensemble diversity showed that the ABB4 models most accurately mirror experimental observations of flexibility. In a secondary analysis, we quantified the coverage of experimentally observed conformational states. While conventional structure prediction models excel at high-resolution predictions of a limited number of states, ABB4-STEROIDS provides a more comprehensive representation of the accessible structural landscape.

We analysed several ablations and model variants to better understand specific contributions to model performance. First, we showed that 50 inference step are required for optimal model performance. Further reductions to 20 or even 10 steps result in minor performance degradation but offer a significant reduction in computational cost. Second, the introduction of a training stage on a large amount of potentially lower quality coarse-grained simulations data prevented overfitting to the MD distributions and enhanced generalisation on the experimental benchmark.

Finally, we evaluated a novel auxiliary loss function specifically targeted at ensemble properties. Generally, structure prediction models, even those designed for conformational sampling, rely on loss functions applied to individual frames. While BioEmu [38] recently introduced a loss calculated over multiple predictions to regress folded versus unfolded proportions, we propose a new loss formulation that minimises the error in residue-level RMSF. Unlike the BioEmu ensemble loss, which requires model rollouts during training and thus incurs significant computational overhead, our loss is integrated into the regular loss computation introducing no additional cost. Predictions for canonical CDRs were substantially improved through the introduction of this loss, however, CDRH3 did not experience a similar performance boost which we attribute to training instabilities caused by the high prediction uncertainty of this loop. Due to the CDRH3 results, we did not include the loss in the final model, nonetheless, findings suggest ensemble-targeted losses may be an effective strategy for optimising structural distributions.

A primary limitation of ABB4-STEROIDS is its exclusive training on unbound antibody simulations and consequently, predictions represent the apo-ensemble. Antigen binding tends to rigidify CDRs [4, 3], therefore, we expect the magnitudes of flexibility to match the unbound and not the bound ensemble. However, as bound conformations are commonly contained in the unbound ensemble [54], our model should capture these states. This hypothesis is supported by good coverage of experimental conformations, most of which are antigen-bound. Modelling the unbound ensemble remains important as it is determines properties critical during drug design such as affinity, specificity and developability. While, predicting conformational changes induced by antigen binding has been attempted [52], modelling the conformational ensembles of the bound state remains an important direction for future work, as it would provide deeper insight into the modulation of functional properties by structural flexibility [9].

In summary, ABB4-STEROIDS is a deep learning model capable of sampling antibody conformational ensembles with high accuracy. The model is released open-source along side the dataset of all-atom antibody simulations used for training. These resources enable improved and large-scale investigations of antibody flexibility with the potential to accelerate rational design of flexibility-modulated properties.

## 4 Methods

### 4.1 Datasets

#### 4.1.1 Single structure prediction dataset

A dataset of experimentally solved structures was created for the single structure prediction task. We used the dataset released with ABB3 [45] and supplemented this set with structures released after its cutoff date.

The ABB3 dataset contains 9042 antibody structures. This corresponds to all experimentally solved structures before 13/12/2023 filtered by the following criteria: resolution better than 3.5 Å, removal of outliers that fall outside of 3.5 standard deviations from the mean for any of the six summary statistics given by ABangle [46], ultra-long CDRH3 loops with more than 30 residues and origin from species which occur less than 15 times in the SAbDAb [48, 49].

To supplement this set, we extracted all antibodies from the SAbDab [48, 49] released between 13/12/2023 and 25/03/2025 and apply the same filtering criteria. We further remove any structures with missing residues between IMGT numbered residues 5 and 120. This resulted in an additional 927 antibody structures, increasing the total size of the dataset to 9969 structures.

#### 4.1.2 Data splits

For evaluation, we created a test set of 100 antibodies with no overlap to the training sets of any method benchmarked in this paper (ABB3, Boltz-1, AF2, aSAM, AlphaFlow and Bioemu). Antibodies in the single structure prediction dataset set released after 31/01/2023, which corresponds to the AlphaFlow [40] training cutoff and is the latest training cutoff of all benchmarked methods, were considered to be included in the test set. Structures with more than 80% identity of the concatenated CDR sequence to any antibody released before the 31/01/2023 with resolution below 9 Å were removed. In this step, we only use a 9 Å resolution cutoff as this is the only filter used in the Boltz-1 training set and corresponds to the most lenient quality filtering by any of the benchmarked methods. In addition, we required test set sequences to have a resolution below 2.5 Å and selected the 100 antibodies with the latest release date as our full test set resulting in a test cutoff on 08/11/23. As ABB3 was not trained with date-based data splits its training set overlaps with our full test set. We, therefore, removed 9 antibodies from the full test set resulting in a reduced dataset of 91 antibodies for comparison against ABB3. A full list of PDB codes of test set antibodies is provided on GitHub (github.com/oxpig/ABB4).

The remaining antibodies in the single structure prediction dataset were clustered at 80% identity of the concatenated CDR sequence using MMseqs2 [55]. 150 antibodies with no overlap to the test set selected as the validation set and the remaining 8205 antibodies with no overlap to test and validation sets were used for training. A total of 1264 antibody structures remained unassigned.

#### 4.1.3 Coarse-grained MD dataset

For ensemble training and evaluation, we created two coarse-grained MD simulation datasets using the Flexible Antibody Database (FlAbDab) [16] and the associated CALVADOS 3-Fv method [16].

We obtained simulations for all non-redundant antibodies in the single structure dataset from FlAbDab. As the database was built using an earlier data cutoff (29/05/2024) than our datasets we used CALVADOS 3-Fv to simulate as many of the remaining antibodies as possible. Antibodies were assigned the same same data split as in as the single structure prediction dataset resulting in 100 test, 150 val and 3187 training simulations. This set contains a much smaller number of training structures as the antibodies were filtered for sequence redundancy which was not performed for the single structure prediction dataset. We refer to this set as high quality simulations.

As part of FlAbDab, a set of ∼ 150, 000 antibody simulations started from predicted structural models was released. A filtered subset of these simulations was used as an additional low quality dataset for model training. Simulations were filtered for by 80% concatenated CDR sequence identity to the test and validation sets as well as outliers in global, heavy chain, light chain, CDRH1, CDRH2, CDRH3 CDRL1, CDRL2 and CDRL3 RMSD and VH-VL distance. An outlier was defined as a deviation by more than 3.5 standard deviations from the mean. This resulted in a dataset of ∼ 136, 000 additional lower quality training simulations.

The coarse-grained MD trajectories in FlAbDab [16] contain 1000 frames. As it is computationally expensive to work with this many frames, we extracted a random subset of 30 frames from each trajectory as a representative ensemble. We make this choice as we observed that RMSF and RMSD values start to converge for most simulations when sampling around 20 frames at random (see SI Section S2.1 for details).

#### 4.1.4 All-atom MD dataset

We simulated a total of 83 antibodies using the protocol outlined in the methods below. Before performing any analysis and further data processing of these simulations the first 60 ns of each replica were discarded. We extracted a representative ensemble of 100 frames for each antibody to create the dataset for model training. Specifically, we sampled 25 frames with a constant time interval of 9.6 ns from each replica.

#### 4.1.5 Datasets of experimental CDR ensembles

To benchmark against experimental data, we created datasets of antibody structural ensembles observed in crystal structures using an approach introduced in [18, 16]. We largely follow the methodology described in [16] but make a small modification to the RMSD calculation.

All antibodies in SAbDab on May 29th 2024 were collected and filtered for structures solved by X-ray crystallography, a resolution lower than 3.5 Å and no missing residues between IMGT numbered positions 5 and 128. Structures were grouped by Fv sequence identity to define an experimental ensemble. For each experimental ensemble the number of observed conformations was calculated for each of the six CDRs individually. Pairwise C*α* RMSD of CDR residues after alignment on C*α* atoms of framework residues of the same chain was calculated (note in [16] structures were aligned on C*α* atoms of CDR residues). A conformation was defined as a cluster resulting from agglomerative clustering with a complete linkage criterion and a RMSD thresholds of 1.25 Å. This approach means that the RMSD of any two structures within a cluster is below 1.25 Å. This threshold was chosen as it was show to result in good functional clustering [56]. Binary labels of flexibility (‘flexible’, ‘rigid’) were assigned to each of the CDRs depending on whether the data suggest a CDR has multiple or a single accessible conformation. CDRs with more than one conformation were labelled as ‘flexible’ and CDRs with a single conformation observed in more than five separate PDB structures were labelled as ‘rigid’. The remaining antibodies were assigned left unlabelled. The final number of data points is shown in Table 1.

**Table 1:** Number of data points in the experimental CDR flexibility datasets.

| DATASET | CDRH1 | CDRH2 | CDRH3 | CDRL1 | CDRL2 | CDRL3 |
| --- | --- | --- | --- | --- | --- | --- |
| TOTAL LABELLED CDRs | 133 | 136 | 220 | 137 | 83 | 110 |
| MULTIPLE CONFORMATIONS | 117 | 117 | 205 | 117 | 48 | 87 |
| SINGLE CONFORMATION | 16 | 19 | 15 | 20 | 35 | 23 |

### 4.2 All-atom MD simulations

To run all-atom MD simulations, we obtained starting structures from the PDB [57] and cleaned each file by removing all non-amino-acid records. We then processed structures with PDBFixer (v1.12) to identify and replace non-standard residues, remove heterogens, model missing residues and add missing heavy atoms, including terminal oxygen atoms at chain C-termini. Before adding hydrogens, we detected disulphide bonds using a combination of automated S*γ*-S*γ* proximity detection and SSBOND record parsing so that cysteines were correctly typed as CYX. We added hydrogen atoms at pH 7.0 using OpenMM Modeller (v8.4.0) [58], with terminal residues handled automatically by the force field template matching.

We parameterised the protein with the AMBER ff19SB force field and used the TIP3P-FB water model [59, 60]. We solvated each system in a truncated octahedral box with a minimum padding of 1.2 nm from the solute to the box edge and set the ionic strength to 0.15 M by adding Na^+^ and Cl^−^ ions. We treated long-range electrostatics using Lennard-Jones particle mesh Ewald with an error tolerance of 5×10^−4^ and a real-space cutoff of 0.9 nm. We applied hydrogen mass repartitioning by redistributing hydrogen masses to 1.5 amu to enable a 4 fs integration time step. We constrained covalent bonds involving hydrogen using SHAKE/SETTLE and kept water molecules rigid. We integrated the equations of motion with the LangevinMiddle integrator using a collision frequency of 1 ps^−1^ and a constraint tolerance of 10^−6^, and we removed centre-of-mass motion at regular intervals.

We performed energy minimisation in three stages using a convergence tolerance of 10 kJ/mol/nm throughout. First, we restrained all non-solvent, non-hydrogen atoms with 100 kcal/mol/Å^2^ for 5,000 iterations. Second, we reduced restraints to 10 kcal/mol/Å^2^ for 2,500 iterations. Third, we removed all restraints and minimised for a further 2,500 iterations. After minimisation, restraint reference positions were updated to the minimised coordinates so that subsequent equilibration restraints pulled toward the relaxed geometry. We equilibrated each system using a nine-stage protocol totalling approximately 7 ns. We assigned velocities from a Maxwell-Boltzmann distribution at 100 K and ran a 20 ps NVT simulation with a 1 fs time step and friction 5 ps^−1^. We then heated the system from 100 K to 298 K over 1 ns under NVT using a linear temperature ramp divided into 200 blocks, using a 2 fs time step, friction 5 ps^−1^, and heavy-atom positional restraints of 50 kcal/mol/Å^2^. We initiated NPT by introducing a Monte Carlo barostat (1 bar, attempt frequency every 25 steps) and ran an initial 50 ps NPT settling period at 2 fs with friction 5 ps^−1^ and a reduced barostat attempt frequency (every 100 steps). We then ran 1 ns of NPT at 4 fs with heavy-atom restraints of 100 kcal/mol/Å^2^, followed by 1 ns of NPT with heavy-atom restraints reduced to 10 kcal/mol/Å^2^. Next, we removed heavy-atom restraints and applied backbone (N, C*α*, C) restraints of 10 kcal/mol/Å^2^, performed an intermediate energy minimisation (5,000 iterations), and continued NPT for 1 ns with backbone restraints of 10 kcal/mol/Å^2^. We then ran successive 1 ns NPT segments with backbone restraints reduced to 1 kcal/mol/Å^2^ and 0.1 kcal/mol/Å^2^, and finished with 1 ns of unrestrained NPT. For each NPT equilibration segment, we used a NaN-recovery protocol where if numerical instabilities occurred, we rolled back to the most recent checkpoint, performed a brief stabilisation with a reduced time step and increased restraints, re-minimised, and retried the segment.

We ran production simulations for 300 ns per replica under NPT conditions (298 K, 1 bar) with all restraints removed and a 4 fs time step, giving a total of 1.2 *µ*s of simulation. We saved trajectory frames every 15,000 steps (60 ps) and recorded thermodynamic observables (potential energy, kinetic energy, total energy, temperature, volume, and density) at the same interval. We wrote checkpoints every 25,000 steps to enable restarts and ran three independent replicas for each antibody structure. We performed all simulations using CUDA-accelerated OpenMM (v8.4.0) with mixed-precision arithmetic on NVIDIA H200 GPUs on the Isambard-AI facility HPCs.

We processed production trajectories using cpptraj in AmberTools24 [61, 62]. For each replica, we used the equilibrated structure as the reference topology, centred trajectories on the protein centre of mass, unwrapped across periodic boundaries, and re-imaged using autoimage. We stripped solvent molecules and ions to generate protein-only trajectories and aligned frames to the first frame by root-mean-square fitting of protein heavy atoms. We performed all trajectory processing on an 8-core CPU allocation with 150 GB memory.

### 4.3 Metrics for ensemble analysis

Metrics were calculated for each antibody region separately. IMGT numbering was used for region definitions [63] and numbering was carried out using the ANARCII software [64].

#### 4.3.1 Evaluation of predicted ensembles against MD data

We used several metrics to compare sampled structures against the MD ensembles. Distance-based metrics recall, precision and distributional accuracy are based on the distance between predicted structures and individual frames from the MD trajectories. We calculated the pairwise RMSD matrix *C* ∈ ℝ^*m*×*n*^ between the *m* structures in the sampled ensemble 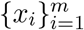 and the *n* representative MD frames 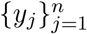.

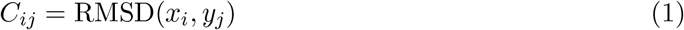

Pairwise RMSD is defined by our standard protocol as the C*α* RMSD of a domain after alignment on all C*α* atoms of the chain the domain is located on. For ‘global’ metrics, structures are aligned on C*α* atoms of both chains. Recall is defined as the mean minimum distance between all MD frames and any sampled structure.

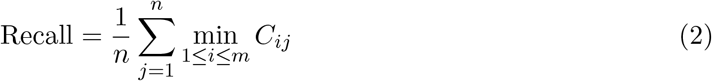

Precision is defined as the mean minimum distance between all sampled structures and any MD frame.

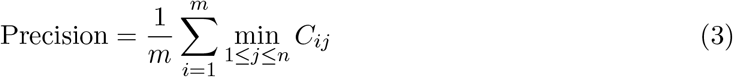

Distributional accuracy is defined as the 1-Wasserstein distance between the distributions of sampled structures and MD frames, using *C* as the ground cost matrix. Assigning uniform weights 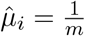 and 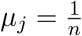, the distance is

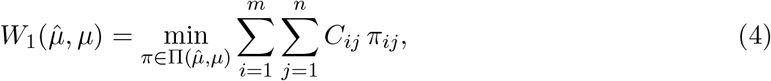

where 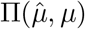 is the set of all couplings with marginals 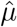 and *µ*.

Ensemble diversity metrics measure the diversity within a predicted ensemble and compare the magnitude of diversity to the MD data. We used the mean absolute error (MAE) of ensemble RMSD and the root mean squared error (RMSE) of ensemble RMSF as two such metrics. To calculate MAE RMSD, we computed the pairwise RMSD matrix *R*^*sampled*^ ∈ ℝ^*m*×*m*^ between the *m* sampled structures of an antibody and the pairwise RMSD matrix *R*^*MD*^ ∈ ℝ^*l*×*l*^ between all *l* frames in the MD trajectories.

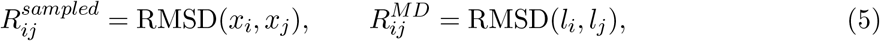

The MAE between the two RMSD matrices over *K* antibodies is defined as

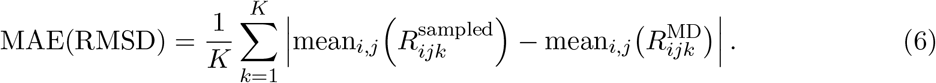

To calculate RMSE of RMSF, we calculate the RMSF of residue *r* in the sampled ensemble 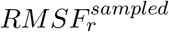 and *l* frames of the MD trajectory 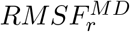. The RMSE of RMSF is defined per antibody as

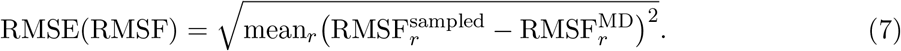

We used time-lagged independent component analysis (tICA) to visually compare the predicted ensemble against the all-atom MD simulations. To perform the tICA, the all-atom MD simulations were subsampled at 240 ps intervals resulting in 4000 frames per antibody. tICA was performed on the *ϕ* and *ψ* backbone torsion angles of each CDR individually.

#### 4.3.2 Evaluation of predicted ensembles against experimental data

We used two metrics, ‘flexibility correlation’ and ‘conformation coverage’, to compare sampled structures against the experimental ensembles.

Flexibility correlation measures the correlation between RMSD in sampled ensembles and flexibility of the experimental ensemble. We calculated the pairwise RMSD matrix *R*^*sampled*^ ∈ ℝ^*m*×*m*^ between the *m* sampled structures of an antibody. For each antibody, we summarized its sampled structural variability by the median pairwise RMSD

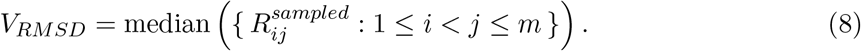

Flexibility correlation is defined as the point biserial correlation coefficient (r) between *V*_*RMSD*_ and the binary flexibility labels of the experimental ensemble, where *Y* = 1 indicates a flexible antibody and *Y* = 0 indicates a rigid antibody. Flexibility correlation was calculated for all antibodies that were labelled as either ‘flexible’ or ‘rigid’.

Conformation coverage measures the coverage of conformations in the experimental ensemble by the sampled structures. We compute the pairwise RMSD matrix *C* ∈ ℝ^*m*×*n*^ between the *m* sampled structures of an antibody and the *n* structures in the experimental ensemble. As a conformational cluster ℰ_*k*_ = {*y*_*j*_ : *j* ∈ *J*_*k*_} may contain more than one structure, we first define its mean distance *D* ∈ ℝ^*m*×*k*^ to a sampled structure *x*_*i*_ as

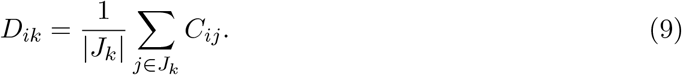

The conformation coverage score is then defined as the minimum distance to any sampled structure

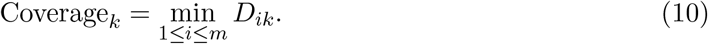

Conformation coverage was only calculated for CDRs with multiple conformations.

### 4.4 ABB4 model

ABB4 is a generative protein structure prediction model using flow matching. The generative process, model architecture and training and inference procedure are briefly outlined below and details provided in the SI.

#### 4.4.1 Model formulation

Protein structure is parametrised, following AlphaFold2 [27], by a backbone rigid-body transformation *T* ∈ *SE*(3) together with a set of side chain torsion angles *χ* for each amino acid residue. We formulate structure prediction as a generative process over backbone frames, modelling a conditional distribution *p*_*θ*_(*T*_1_ | *T*_0_, *s*), which interpolates between an initial noisy frames *T*_0_ and the final protein backbone configuration *T*_1_ conditioned on the sequence *s*. Torsion angles are predicted via *p*_*θ*_(*χ* | *T*_1_), conditioned on the final backbone frame.

Backbone frames *T* = (*x, r*) consist of a translation vectors *x* ∈ ℝ^3×*N*^ and a rotation matrices *r* ∈ *SO*(3)^*N*^. We modelled the generative process using the flow-matching formulation on *SE*(3) introduced in FrameFlow [44]. In brief, translations and rotations are denoised independently through a learned vector field *p*_*θ*_(*v*_*t*_ | *T*_*t*_, *t, s*), where *v* = (*v*_*x*_, *v*_*r*_). In practice, the model is parametrised to predict clean frames 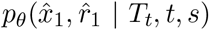, from which we construct the corresponding vector field and integrate it to obtain the next sample in the trajectory 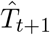,

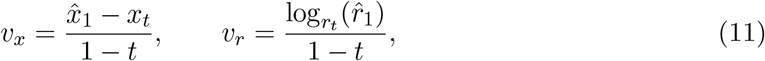

where 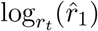 denotes the logarithmic map on *SO*(3), corresponding to the geodesic direction from *r*_*t*_ to 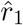. For further details on flow matching in *SE*(3) see FrameFlow [44].

The model is trained with direct losses on the backbone translations ℒ_*R*(3)_ and rotations ℒ_*SO*(3)_ as in FrameFlow [44] and auxiliary FAPE ℒ_*FAPE*_, torsion angle ℒ_*χ*_ and torsion norm ℒ_∥*χ*∥_ losses on all-atom coordinates and torsion angle predictions as defined as in AlphaFold2 [27]. Details on individual loss terms are provided in SI Section S1.1.1. The total loss is defined as:

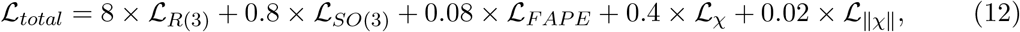

with weights set to match the magnitude of individual terms after losses plateaued. All of these losses have the objective to reconstruct a single structure.

We additionally implement a loss that acts across an ensemble of generated structures ℒ_*ensemble*_ with the objective to fit the RMSF to the expectation in the MD simulations (for details see SI Section S1.1.2). The main ABB4 models do not use the ensemble loss but we evaluated a model variant trained with this loss. For this model the ensemble loss it is added to the total loss with a weight of 0.05.

We used a global sequence-conditioned optimal transport strategy for training. In contrast to previous work, which employed unconditional, mini-batch optimal transport [65], we implemented a variant tailored to the task of ensemble prediction. For each protein sequence s, we compute a global optimal transport coupling 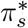 over all noise-structure pairs associated with that sequence. It should be noted that while the structural frames remain constant throughout training, the Gaussian noise is resampled at every epoch to ensure robust performance. Accordingly, we optimize

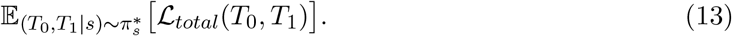

This approach preserves the benefits of optimal transport while conditioning on biological sequence identity, which ensures consistency across all conformations of a given protein sequence. For more details see SI Section S1.1.3.

#### 4.4.2 Model architecture

The ABB4 model builds on invariant point attention (IPA) first introduced in AlphaFold2 [27] and largely follows the architecture introduced in FrameDiff [66].

The model takes as input: node features *H*_0_, edge features *Z*_0_ and initial backbone frames *T*_0_. The node features are constructed from an amino acid embedding, a positional embedding, a time-step embedding and an antibody region embedding. The antibody region embedding is a 14-dimensional one-hot vector indicating which region (CDRH1, CDRH2, CDRH3, CDRL1, CDRL2, CDRL3, FwH1, FwH2, FwH3, FwH4, FwL1, FwL2, FwL3, FwL4) the residue belongs to. For an antibody of *r* residues, these are embedded into an initial node feature encoding *H*_0_ ∈ ℝ^*r*×256^. Edges encode a distogram of backbone frame translations and a relative positional encoding. These are embedded into an initial edge feature encoding *Z*_0_ ∈ ℝ^*r*×*r*×256^.

Node features, edge features and backbone frames are updated by 8 layers of a structure module (Figure 7). The final layer of the model outputs the backbone rotations and translation as well as side chain torsion angles which are used to reconstruct an all-atom structure. ABB4 has a total of 19.6 million trainable parameters.

**Figure 7:**
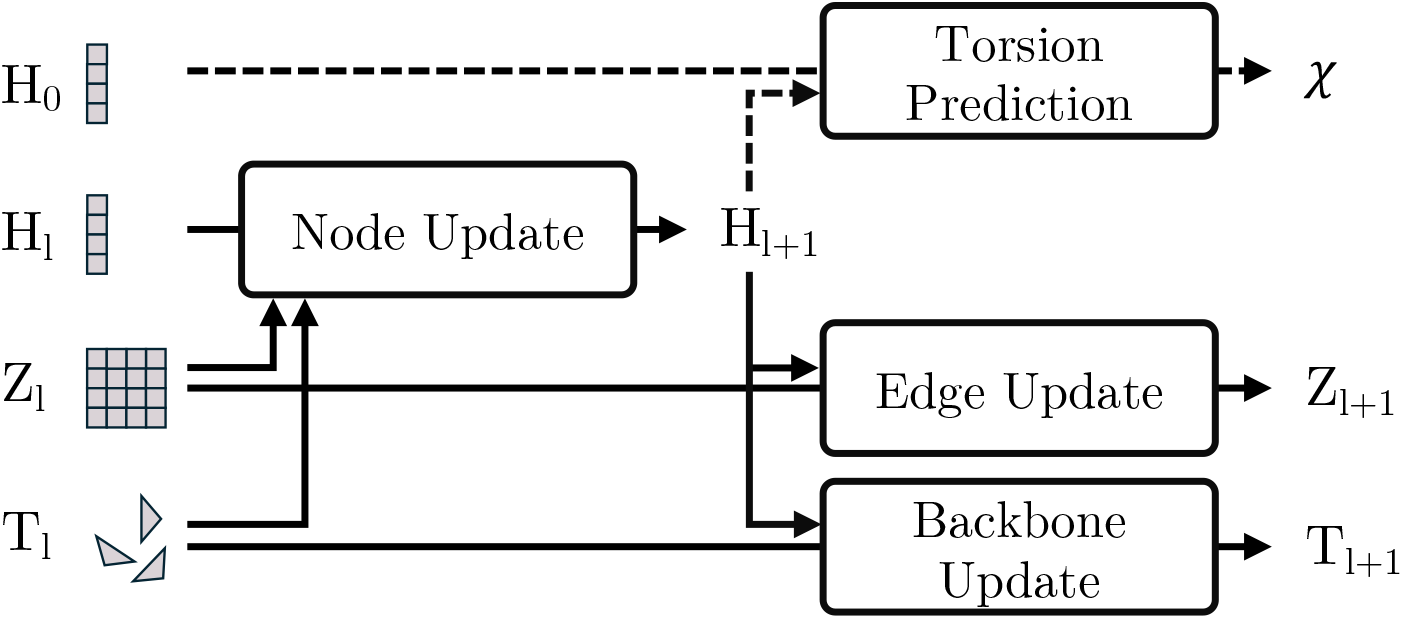
Single layer of the ABB4 model. Each layer *l* has the current node representations *H*_*l*_, edge representations *Z*_*l*_ and backbone frames *T*_*l*_ and initial node representations *H*_0_. Representations are processed with node, edge and backbone update blocks to prepare the input for layer *l* + 1. In the final layer (indicated by the dashed line), torsion angles *χ* are additionally predicted. Details on the architecture of the four model blocks are shown in Figure S4.

#### 4.4.3 Training

We trained three variants of the ABB4 model, which we refer to as ABB4-base, ABB4-STEROIDS-CG, and ABB4-STEROIDS. These differ by the set of training stages applied. In total, we used four distinct stages to train the models. During stage 1, we trained on on the single structure prediction dataset for the task of single structure prediction. We refer to the resulting model as ABB4-base. In the following two stages, the base model was trained on representative ensembles from the coarse-grained MD dataset for the task of ensemble prediction. In stage 2, we trained on all available simulation including the high quality set and the 136,000 lower quality simulations started from predicted models. In stage 3, only the high quality set of simulations was used. We refer to the model trained with stages 1 to 3 as ABB4-STEROIDS-CG. In stage 4, we further fine-tune the model on a smaller amount of all-atom MD simulation and obtain the final ABB4-STEROIDS model. The total dataset sizes used in each stage are shown in Table 2.

**Table 2:** Training dataset sizes.

| STAGE | DATASET | TRAINING |  | VALIDATON |  | TEST |  |
| --- | --- | --- | --- | --- | --- | --- | --- |
|  |  | MABS | STRUC. | MABS | STRUC. | MABS | STRUC. |
| 1 | SINGLE STRUCTURE PRED. | 8205 | 8205 | 139 | 139 | 100 | 100 |
| 2 | CG MD ALL | 136,000 | 4,180,000 | 1480 | 4440 | 100 | 3000 |
| 3 | CG MD HIGH QUALITY | 3187 | 95,000 | 1480 | 4440 | 100 | 3000 |
| 4 | ALL-ATOM MD | 73 | 7150 | 4 | 400 | 14 | 1400 |

mAbs: number of unique antibodies, Strucs: total number of crystal structures/MD frames, CG: coarse-grained.

Models were trained with a batch size of 32 (stage 1) and 40 (stage 2, 3 & 4). The AdamW optimiser was used with weight decay of 0.01. A cosine annealing with warm restarts learning rate schedule was used with an initial learning rate of 1e-4, a minimum learning rate of 1e-5 and 50 epochs interval between restarts.

We used early stopping based on validation set metrics. During stage 1, we performed a mini roll-out with 10 model evaluations to generate one structure for each validation set antibody and compute the RMSD to the crystal structure. We stopped training when the CDRH3 RMSD converged. During stage 2, 3 and 4, we roll-out an ensemble of 30 structures for each validation set antibody and compute RMSD within the generated ensemble. We stopped model training when the error in RMSD between the generated ensemble compared to the full MD trajectory converged for CDRH1 (stage 2) and CDRH3 (stage 3 & 4). We found this approach using mini roll-outs results in much improved performance over monitoring of step-level validation losses.

#### 4.4.4 Inference

Unless stated otherwise, inference was performed with 100 model evaluations. An ensemble of 100 structures was sampled for each antibody during evaluation.

### 4.5 Inference of benchmarked models

In this section, details on how inference was run with any of the benchmarked models is provided. When evaluating a model for single structure prediction (ABB3, Boltz-1), a single structure was generated for each antibody. For evaluation of ensemble prediction (Boltz-1, BioEmu, AlphaFlow, aSAM, AF2 MSA subsampling), 100 structures were sampled for each antibody.

ABB3, Boltz-1 and AF2 allow for inputs with multiple chains and can, therefore, be used directly to predict antibody structures. In contrast, BioEmu, AlphaFlow and aSAM accept inputs with only a single chain. For these methods, we place heavy and light chain sequences on a single chain connected by six repeats of a glycine and serine linker (6 × *GGGGS*).

#### 4.5.1 ABB3

ABB3 inference was performed according to authors instruction [45]. Input embeddings were produced using the default T5 pLM model distributed with the package.

#### 4.5.2 Boltz-1

Inference was performed with Boltz-1 weights[37] and default parameters. MSAs were generated through the Boltz-1 using the MMseqs2. For single structure prediction we set the number of diffusion sampled to 1 and for ensemble prediction to 100.

#### 4.5.3 BioEmu

Backbone structures were sampled from the BioEmu [38] model with default parameters. Side chains were reconstructed using the provided script. MD equilibration was not performed.

#### 4.5.4 AlphaFlow

The AlphaFlow-MD distilled model with default parameters was used for inference [40]. The distilled model was chosen over the base model to reduce the computational cost. MSAs were generated using the provided script to query the ColabFold [67] servers.

#### 4.5.5 aSAM

aSAM [39] does not perform structure prediction and requires a starting structure as input. Three structures of each antibody were generated with Boltz-1 and selected the highest ranked structure as the input structure for aSAM. We used the ATLAS model for inference with default parameters.

#### 4.5.6 AlphaFold2 MSA subsampling

AF2 was run through ColabFold [67]. For MSA subsampling, the max MSA parameter was set to 64:128, the number of recycles to 1 and the number of seeds to 20 (5 models are generated per seed which results in 100 total structures). These parameters have previously been shown to generate antibody ensembles with maximal diversity while limiting the occurrence of unfolded structures [18].

## Data availability

The all-atom MD dataset is released on Zenodo (https://doi.org/10.5281/zenodo.19471889).

## Code availability

All ABB4 models (github.com/oxpig/ABB4) and scripts used to perform the all-atom antibody MD simulations (github.com/oxpig/antibodymd) are openly available on GitHub.

## Acknowledgements

This work was supported through research funding from the UK Engineering and Physical Sciences Research Council (EPSCR) [Grant number EP/S024093/1], Roche and the Royal Commission for the Exhibition of 1851 awarded to FS. The authors acknowledge the use of resources provided by the Isambard-AI National AI Research Resource (AIRR). Isambard-AI is operated by the University of Bristol and is funded by the UK Government’s Department for Science, Innovation and Technology (DSIT) via UK Research and Innovation; and the Science and Technology Facilities Council [ST/AIRR/I-A-I/1023].

## Author contributions statement

FS, MC and CD contributed to the conception and design of the study. FS and OV developed the ABB4 code base. FS, MC, and KI created the datasets. FS trained the models, performed model benchmarking and wrote the manuscript. FS, MC, KI, OV and CD contributed to critical revision of the manuscript. All authors approved the submitted version.

