## Supplementary Information for "Sampling antibody conformational ensembles with ABodyBuilder4-STEROIDS"

### Supplementary Information
### Sampling antibody conformational ensembles with ABodyBuilder4-STEROIDS

Fabian C. Spoendlin[1], Matteo Cagiada[1,2], King Ifashe[1], Odysseas Vavourakis[1], and Charlotte M. Deane[1,*]

[1]Department of Statistics, University of Oxford, Oxford, UK
[2]Department of Biology, University of Copenhagen, Copenhagen, Denmark

**S3 Supplementary figures** **8**

### S1 Supplementary methods

#### S1.1 ABB4 model details

##### S1.1.1 Loss functions

ABB4 is trained using a loss composed of five terms. The model predicts clean backbone frames $\hat{T}_1 = (\hat{x}_1, \hat{r_1})$ and torsion angles $\hat{\chi}$, which together are used to reconstruct all-atom coordinates $\hat{a}$. We optimise the model under an $SE(3)$ flow matching objective following the formulation of FrameFlow (Yim et al., 2023), using separate losses on the translational and rotational components in $R(3)$ and $SO(3)$, respectively:

$$\mathcal{L}_{R(3)} = \mathbb{E}_{(T_0,T_1|s)\sim\pi_s^*,\, s\sim\mathcal{D},\, t\sim\mathrm{U}(0,1)}\left[\frac{1}{(1-t)^2}\sum_{i=1}^{N_{frames}}\left(\left\|\hat{x}_1^i(T_t,t,s) - x_1^i\right\|_{\mathbb{R}^3}^2\right)\right], \quad (1)$$

$$\mathcal{L}_{SO(3)} = \mathbb{E}_{(T_0,T_1|s)\sim\pi_s^*,\, s\sim\mathcal{D},\, t\sim\mathrm{U}(0,1)}\left[\frac{1}{(1-t)^2}\sum_{n=1}^{N_{frames}}\left(\left\|\log_{r_t^i}\left(\hat{r}_1^i(T_t,t,s)\right) - \log_{r_t^i}\left(r_1^i\right)\right\|_{SO(3)}^2\right)\right], \quad (2)$$

where $\pi_s^*$ denotes the optimal-transport coupling (for details see SI Section S1.1.3), $U(0,1)$ denotes the uniform distribution over time and $log_{r_t}(r_1)$ is the Riemannian logarithm on $SO(3)$, corresponding to the tangent-space geodesic direction at $r_t$ pointing toward $\hat{r}_1$.

To stabilise the flow over backbone frames and to introduce side chain prediction capabilities, we include auxiliary losses on the reconstructed all-atom coordinates $\hat{a}$ and torsion angles $\hat{\chi}$. First, we use the frame-aligned point error (FAPE) (Jumper et al., 2021):

$$\mathcal{L}_{\text{FAPE}} = \mathbb{E}_{(T_0,T_1|s)\sim\pi_s^*,\, s\sim\mathcal{D},\, t\sim\mathrm{U}(0,1)}\left[\frac{1}{N_{\text{frames}}\times N_{\text{atoms}}}\sum_{i=1}^{N_{\text{frames}}}\sum_{j=1}^{N_{\text{atoms}}}\right.$$
$$\left.\min\left(\left\|\hat{T}_1^{i^{-1}}(T_t,t,s)\circ\hat{a}_1^j(T_t,t,s) - T_1^{i^{-1}}\circ a_1^j\right\|_{\mathbb{R}^3}^2,\ d_{\text{clamp}}\right)\right], \quad (3)$$

where $T^{-1}$ denotes the inverse rigid transformation and $d_{clamp}$ clamps the deviation at 10Å.

Following AlphaFold2 (Jumper et al., 2021), torsion angles $\chi$ are parametrised as points on the unit circle, $\chi_n^i \in \mathbb{R}^2$ with $\|\chi^{i,n}\| = 1$. The torsion parametrisation exhibits a 180°-rotation-symmetry, meaning that each torsion angle corresponds to two equivalent unit-circle vectors. AlphaFold2 supervises the prediction against either of these two symmetric targets. We simplify the torsion loss by supervising only a single representative orientation, which we find does not affect performance.

The torsion angle loss is

$$\mathcal{L}_{\chi} = \mathbb{E}_{(T_0,T_1|s)\sim\pi_s^*,\, s\sim\mathcal{D},\, t\sim\mathrm{U}(0,1)}\left[\frac{1}{N_{\text{frames}} \times N_{\text{torsions}}} \sum_{i=1}^{N_{\text{frames}}} \sum_{n=1}^{N_{\text{torsions}}} \left( \left\| \frac{\hat{\chi}^{i,n}(T_t,t,s)}{\|\hat{\chi}^{i,n}(T_t,t,s)\|} - \chi^{i,n} \right\|^2 \right) \right], \tag{4}$$

and we additionally regularise the predicted torsion-vector norm to encourage predictions to remain on the unit circle:

$$\mathcal{L}_{\|\chi\|} = \mathbb{E}_{(T_0,T_1|s)\sim\pi_s^*,\, s\sim\mathcal{D},\, t\sim\mathrm{U}(0,1)}\left[\frac{1}{N_{\text{frames}} \times N_{\text{torsions}}} \sum_{i=1}^{N_{\text{frames}}} \sum_{n=1}^{N_{\text{torsions}}} \left( \left\| \|\hat{\chi}^{i,n}(T_t,t,s)\| - 1 \right\| \right) \right]. \tag{5}$$

##### S1.1.2 RMSF ensemble loss

In addition to per-structure losses, we introduced an ensemble loss with the objective to match the RMSF of ensembles generated by our model to the RMSF observed in MD simulation directly. Specifically, for each protein sequence s, we predict an ensemble $\hat{E} = \{\hat{T}_1^1, \ldots, \hat{T}_1^K\}$ of $K$ structure and compute RMSF of $C_\alpha$ atoms across the ensemble. We then penalise the deviation between the predicted RMSF and the reference (MD) RMSF using a normalized $\chi^2$-style loss. This gives the objective

$$\hat{E}(E_T,t,s) = \{\hat{x}_1^1(T_t^1,t,s)), \ldots, \hat{x}_1^K(T_t^K,t,s))\}, \tag{6}$$

$$\mathcal{L}_{ensemble} = \mathbb{E}_{(E_0,E_1|s)\sim\pi_s^*,\, s\sim\mathcal{D},\, t\sim\mathrm{U}(0,1)}\left[\frac{1}{N_{\text{frames}}} \sum_{i=1}^{N_{\text{frames}}} \left( \frac{\left\| RMSF^i(\hat{E}(E_T,t,s)) - RMSF^i(E) \right\|^2}{(\sigma^i)^2} \right) \right], \tag{7}$$

where $x$ is the backbone frame translation which corresponds to the $C_{alpha}$ coordinate, $\mathrm{RMSF}^i(\cdot)$ denotes the root-mean-square fluctuation of residue $i$ across the ensemble, and $(\sigma^i)^2$ is a per-residue variance normalisation term. In our experiments we set $\sigma^i = 0.1 \times \mathrm{RMSF}^i$. Using a $\chi^2$-style (variance-normalised) loss rather than a standard MSE stabilises training by accounting for heteroscedastic fluctuation amplitudes across different residues, so that highly flexible regions do not dominate the loss.

##### S1.1.3 Global sequence-conditional optimal transport

In our setting, we have two probability distributions of noise $p(T_0)$ and protein structures $p(T_1)$. The goal of optimal transport (OT) is to move mass according to $p(T_0)$ so that it becomes distributed as $p(T_1)$, while minimising a given cost of movement $c(x,y)$. A transport plan $\pi$ is a joint distribution over $(x,y)$ with marginals $p(T_0)$ and $p(T_1)$ and we denote the set of all such couplings by $\Pi(p(T_0), p(T_1))$. Formally, the OT problem is given by (Villani, 2009):

$$\pi^* = \arg \min_{\pi \in \Pi(p(T_0), p(T_1))} \mathbb{E}_{(x,y)\sim\pi}\big[c(x,y)\big]. \tag{8}$$

By constructing simpler integration paths, optimal transport leads to faster and more stable training and was shown to improve performance of certain tasks (Bose et al., 2023). In recent protein structure design models, OT couplings were computed unconditionally $(p(T_0), p(T_1)) \sim \pi^*$ and couplings were approximated over a mini-batch to limit the computational cost Bose et al. (2023); Yim et al. (2023). In protein structure prediction, structures are sampled conditional on a sequence $p(T_1 \mid s)$ in contrast to protein design where we sample from an unconditional distribution $p(T_1)$. We therefore construct a sequence-conditional optimal coupling $(p(T_0), p(T_1) \mid s) \sim \pi_s^*$. As the sequence conditionality reduces the number of samples over which the coupling is calculated, we can determine the exact global OT rather than approximating over a mini-batch. Importantly, this computation can be done outside the training loop, so it does not add overhead during training.

Concretely: let $(T_1^1, \ldots, T_1^K)$ denote all available structures (conformations) for sequence s. We sample translation noise $x_0^k \sim \mathcal{N}(0, I_3)^N$ and rotation noise $r_0^k \sim \mathcal{U}(SO3)^N$, then compute the cost matrix $c^{i,j}(T_0^i, T_1^j)$ using Euclidean distance after alignment in $R(3)$

$$x^i_{0,aligned} = align(x_0^i, x_1^i) \tag{9}$$

$$c^{i,j}_{R(3)} = \|x_1^i - x^j_{0,aligned}\|, \tag{10}$$

and the relative angle in $SO(3)$

$$r^{i,j}_{rel} = r_1^i (r_0^j)^T \tag{11}$$

$$c^{i,j}_{SO(3)} = \arccos\Big(\frac{\mathrm{Tr}(r^{i,j}_{\mathrm{rel}}) - 1}{2}\Big), \tag{12}$$

summing to a total cost of

$$c^{i,j}_{SE(3)} = c^{i,j}_{R(3)} + 10 \times c^{i,j}_{S0(3)}, \tag{13}$$

with weights chosen to balance the magnitude of both terms (Bose et al., 2023).

By contrast to FoldFlow and FrameFlow, which sample initial rotations from the isotropic-Gaussian noise distribution ($r_0^i \sim \mathcal{IG}(SO3)^N$), we sample from a uniform distribution ($r_0^i \sim \mathcal{U}(SO3)^N$). In FrameFlow, sampling from $\mathcal{IG}(SO3)^N$ was described to improve performance. We speculate that this improvement arises because sampling from $\mathcal{IG}(SO3)^N$ effectively ensures that rotational noise is always optimally coupled. As FoldFlow only computes OT coupling over $R(3)$ explicitly, this sampling results in a separate OT coupling over $SE(3)$. This may lead to undesirable behaviour during inference when rotation and translation noise is sampled that was coupled to different output structures during training. Under the formulation to perform OT jointly over $SE(3)$, sampling from $\mathcal{IG}(SO3)^N$ or $\mathcal{U}(SO3)^N$ is equivalent.

#### S2 Supplementary results

##### S2.1 Sampling of coarse-grained MD frames.

Simulations in FlAbDab produced with CV3-Fv (Cagiada et al., 2025) consist of 1000 trajectory frames. As it is computationally expensive to work with such a large number of frames, we used set of representative frames for model training and parts of the analysis. To determine the optimal number of frames to use as representatives, we computed how well randomly sampled ensembles of $N$ frames reproduce ensemble metrics calculated across the entire trajectory. We observed that the error in RMSD and RMSF, when computed from randomly sampled frames compared to values across the entire trajectory, start to converge at an ensemble size of $N = 20$ (Figure S1 & S2). To leave a margin for errors, we chose to sample 30 frames from each trajectory as a representative ensemble.

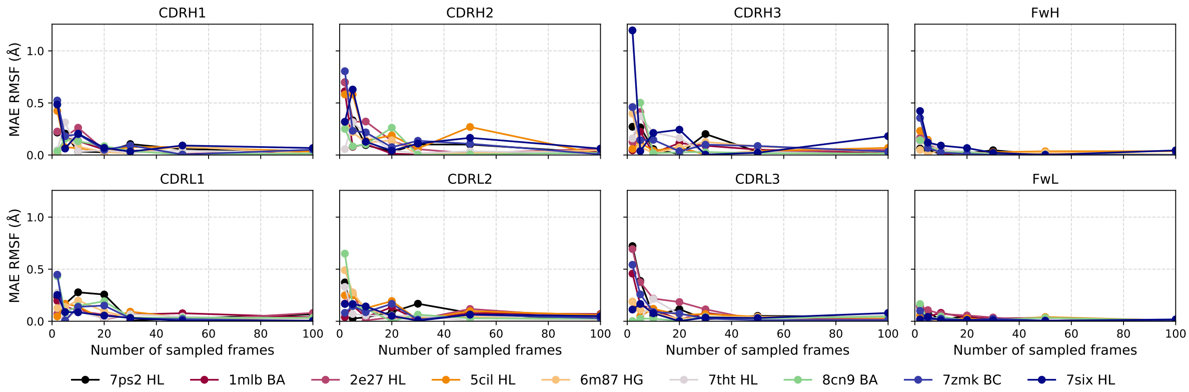

Figure S1: Error in RMSD of sampled MD trajectory frames. The mean absolute error in RMSD of an ensemble of sampled frames compared to the entire MD trajectory is plotted against the number of frames sampled for each ensemble. Values are shown for 9 randomly sampled antibody systems.

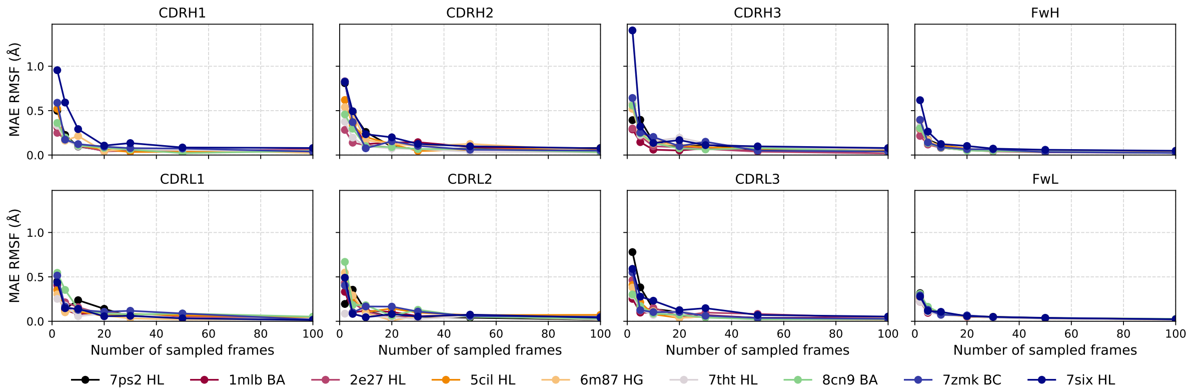

Figure S2: Error in RMSF of sampled MD trajectory frames. The mean absolute error in RMSF of an ensemble of sampled frames compared to the entire MD trajectory is plotted against the number of frames sampled for each ensemble. Values are shown for 9 randomly sampled antibody systems.

#### S2.2 Increased CDRH3 flexibility in FlAbDab simulations started from predicted structural models

FlAbDab (Cagiada et al., 2025) contains a set of simulations started from experimental simulations and a much larger set of simulations started from predicted structural models with ABodyBuilder2 (Abanades et al., 2023). To assess the quality of these simulations, we compared the average RMSF and RMSD of the two set (Figure S3 and observe that the flexibility of CDRH3s in simulations started from predicted structural models tends to be larger. This could be linked to findings that ABB2 predicts CDRH3s with increased solvent exposure (Raybould et al., 2019) which might in turn lead to reduced steric constraints and increased flexibility in simulations.

When training ABB4-STEROIDS in stage 2, we observed that a reduction in the training loss does not correspond to a better reproduction of CDRH3 RMSD and RMSF values evaluated on the validation and test sets. We linked this finding to the increased flexibility of CDRH3s of antibodies in the stage 2 training set (mainly simulations started from predicted models) compared to the validation and test set (only containing high quality simulations started from experimental structures).

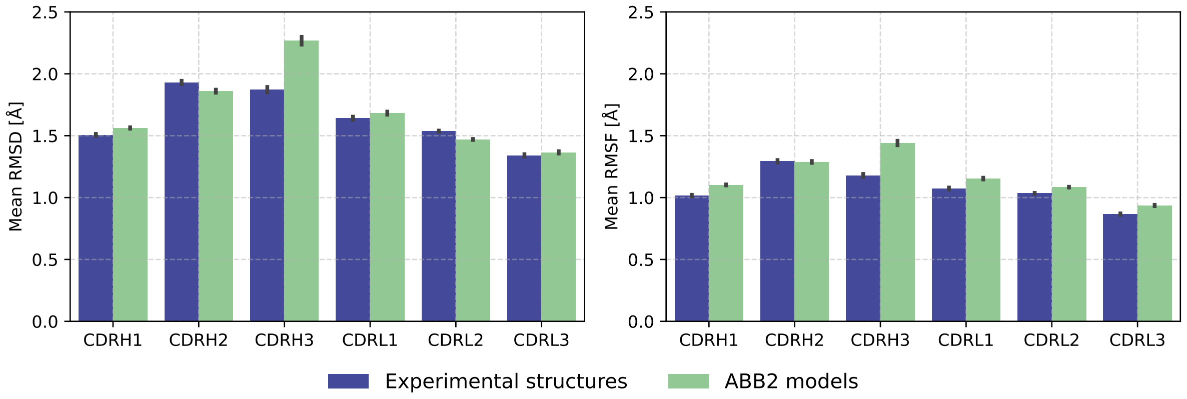

Figure S3: Comparison of CDR flexibility in FlAbDab simulations started from experimental and predicted structural models. Flexibility was assessed in all 4220 simulations started from experimental structures and a sample of 4220 randomly sampled simulations started from structural models. Comparison of mean RMSD values (left) and mean RMSF values (right) for all CDR loops.

#### S3 Supplementary figures

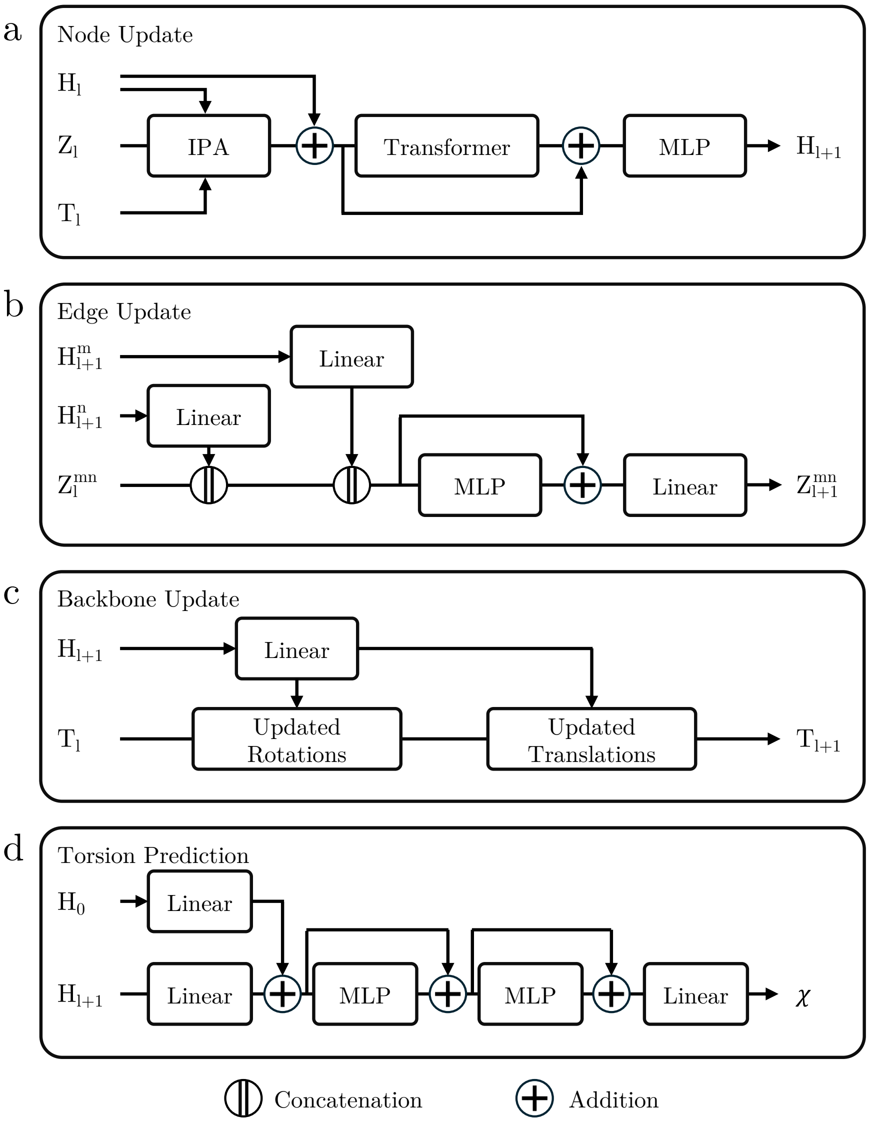

Figure S4: Detailed architecture of the ABB4 models. ABB4 consists of four main blocks: a node update block (a), an edge update block (b), a backbone update block (c) and a torsion prediction block (d). Node features $H$, edge features $Z$ and backbone frames $T$ get updated and torsion angles $\chi$ predicted in the corresponding blocks. Subscript denote the layer $l$ of the model and superscripts the residue indices. IPA indicates an invariant point attention module, MLP a multilayer perceptron and Transformer a transformer encoder module. Updated translation and rotations indicate a direct operation on the translations vector and rotations matrix.

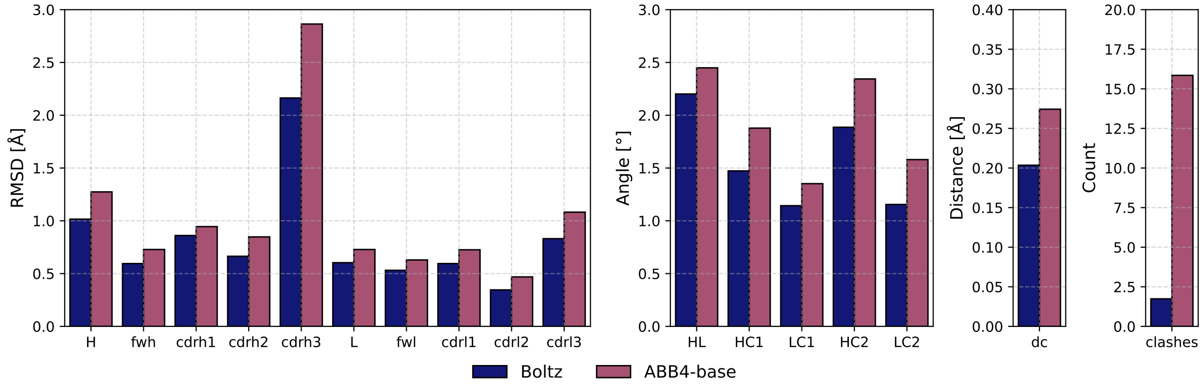

Figure S5: Evaluation of ABB4-base single structure prediction against Boltz-1. Single structure prediction was evaluated on a test set of 100 antibody structures. Predictions were assessed using RMSD of antibody regions (H: heavy chain, fwh: heavy chain framework, L: light chain, fwl: light chain frame work), deviations in VH-VL orientation angels (HL, HC1, LC1, HC2, LC2) and distance (dc) as defined in (Dunbar et al., 2013) and the number of atomic clashes defined as an overlap in van der Waals radii of any backbone or side chain atoms.

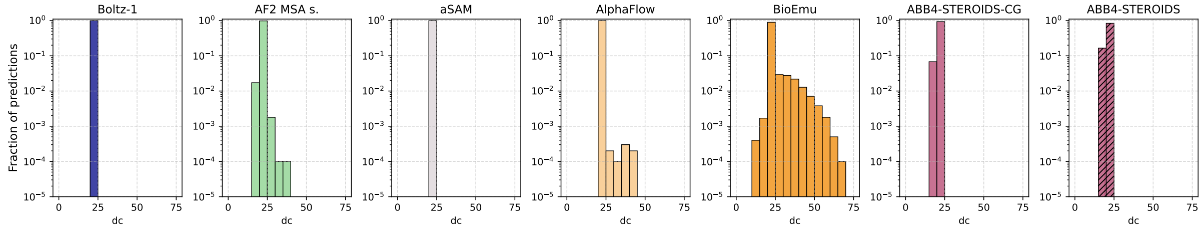

Figure S6: Dissociation of VH and VL domains in predictions. Distributions of VH-VL distance (dc), defined as the distance between the centre of mass of VH C$\alpha$ atoms and VL $C\alpha$ atoms). A distance larger than 30 is indicative of the VH and VL domains dissociating. Fractions are reported from a total of 100 prediction of the full test set of 100 antibodies (10,000 models in total).

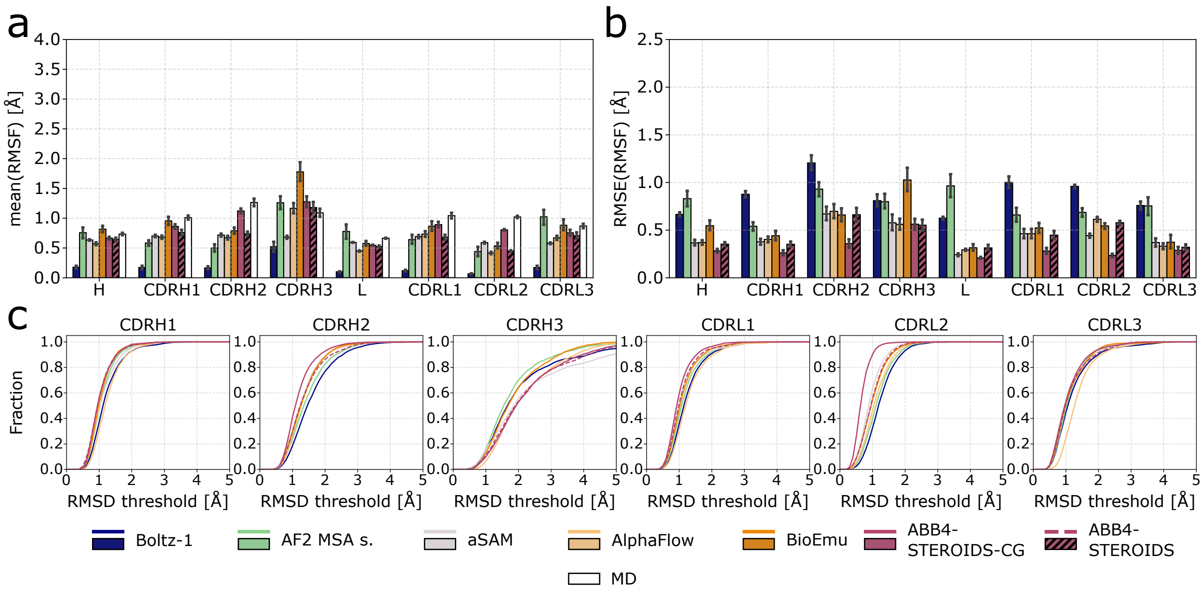

Figure S7: Extended evaluation of predicted structural ensembles against coarse-grained MD data. a) Mean ensemble RMSF in predictions and MD trajectories across all antibodies in the coarse-grained test set. Values computed across the heavy chain (H), light chain (L) and individual CDRs. b) Root mean square error (RMSE) of ensemble RMSF in the predictions compared to the coarse-grained MD data. c) Coverage of MD conformational frames as a function of distance threshold.

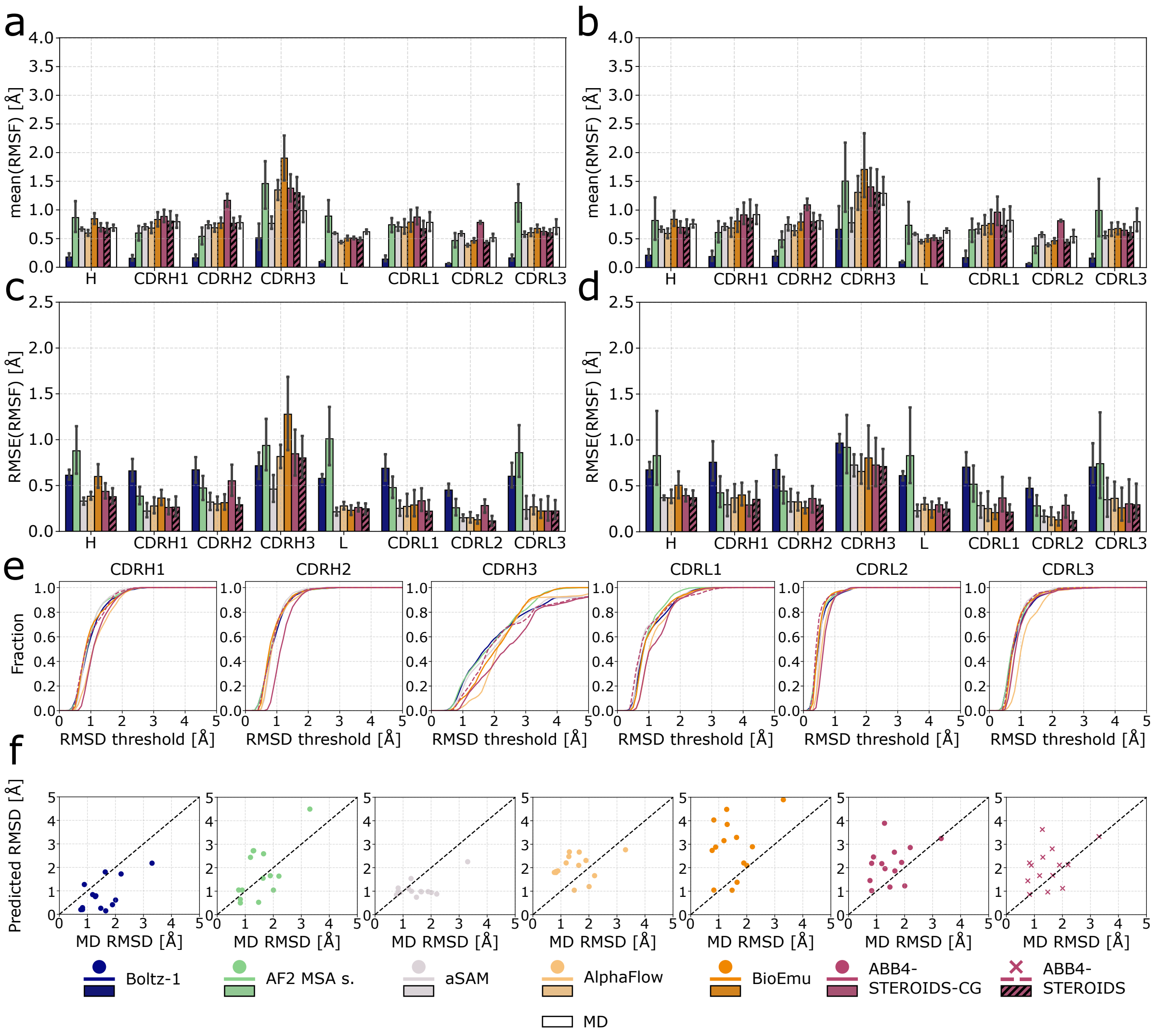

Figure S8: Extended evaluation of predicted structural ensembles against all-atom MD data. Mean ensemble RMSF in predictions and MD trajectories (a) across all antibodies in the all-atom test set and (b) across antibodies with flexible CDRH3s (ensemble RMSD > 1.4 Å in MD trajectories). Values computed across the heavy chain (H), light chain (L) and individual CDRs. Root mean square error (RMSE) of ensemble RMSF in the predictions compared to the all-atom MD data for c) all antibodies in the test set and (d) antibodies with flexible CDRH3s. e) Coverage of MD conformational frames as a function of distance threshold. f) Scatter plot of CDRH3 RMSD in model predictions against CDRH3 RMSD in the all-atom MD. The dashed black line indicates the diagonal corresponding to perfect model predictions.

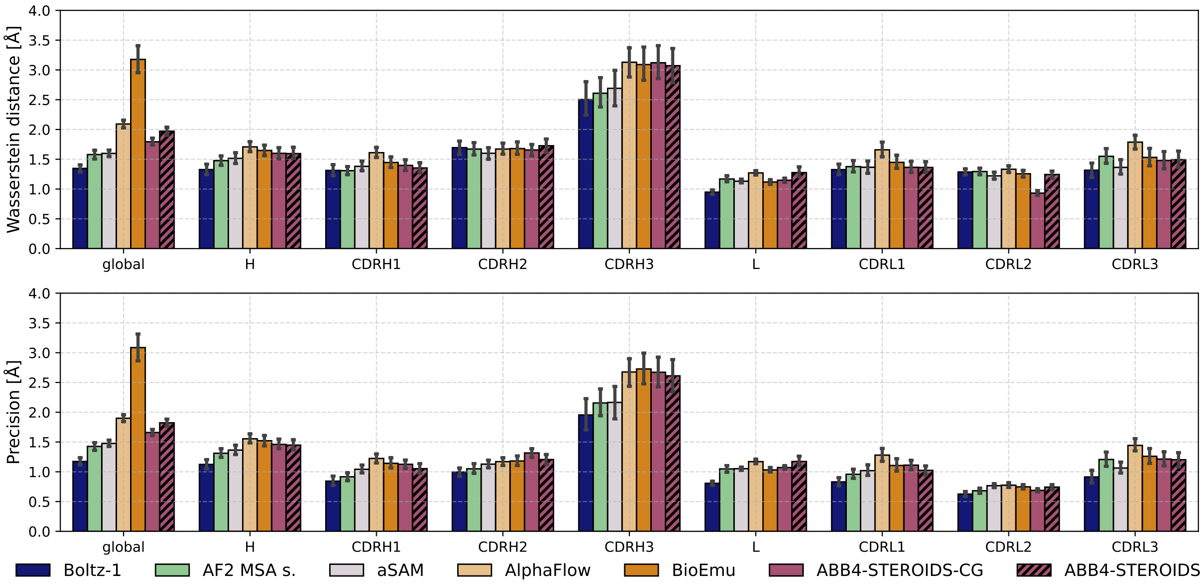

Figure S9: Evaluation of predicted structural ensembles against MD trajectories using distance-based metrics. Wasserstein distance (a) and precision (b) between predicted ensembles and representative frames from MD simulations. Values computed across the entire antibody (global), heavy chain (H), light chain (L) and individual CDRs.

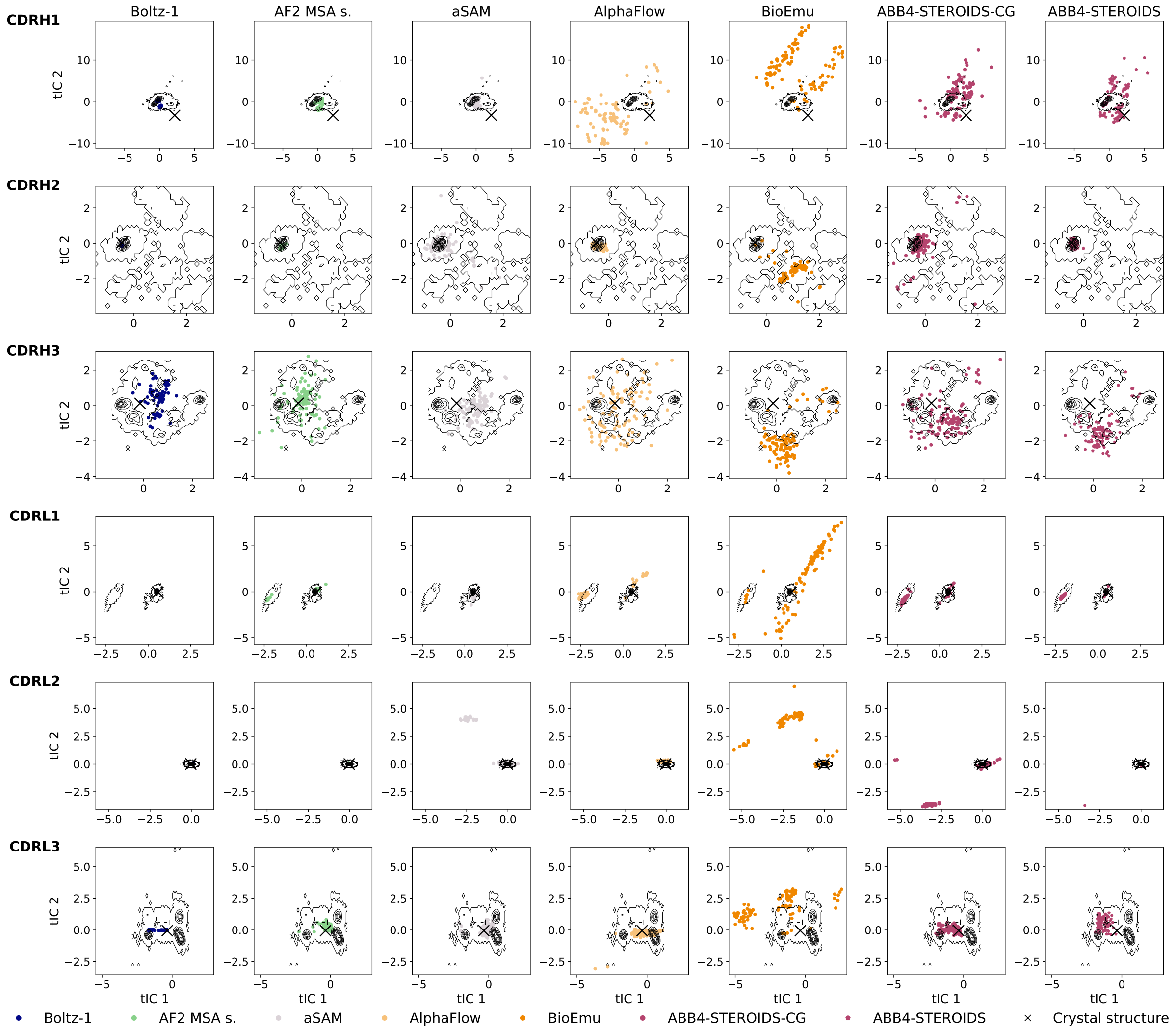

Figure S10: Comparison of the conformational ensemble sampled for test set antibody PDB 8d01. The first two time-lagged independent components (tICs) are plotted. The black contour indicates the all atom MD distributions. Predicted structures were mapped to the tICs and are represented by the coloured dots. The crystal structure from which the simulations was started is shown by the black cross.

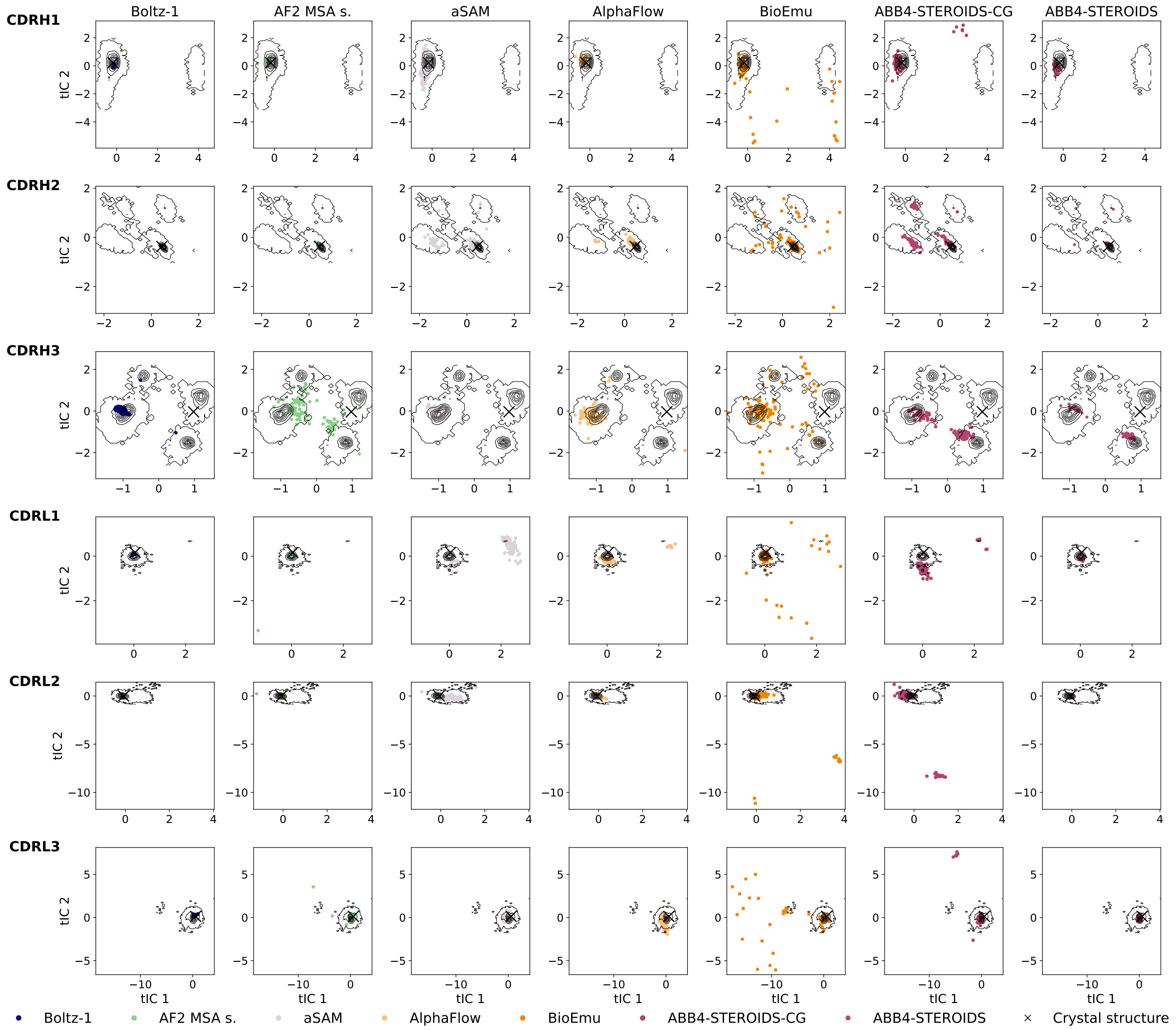

Figure S11: Comparison of the conformational ensemble sampled for test set antibody PDB 8f2t. The first two time-lagged independent components (tICs) are plotted. The black contour indicates the all atom MD distributions. Predicted structures were mapped to the tICs and are represented by the coloured dots. The crystal structure from which the simulations was started is shown by the black cross

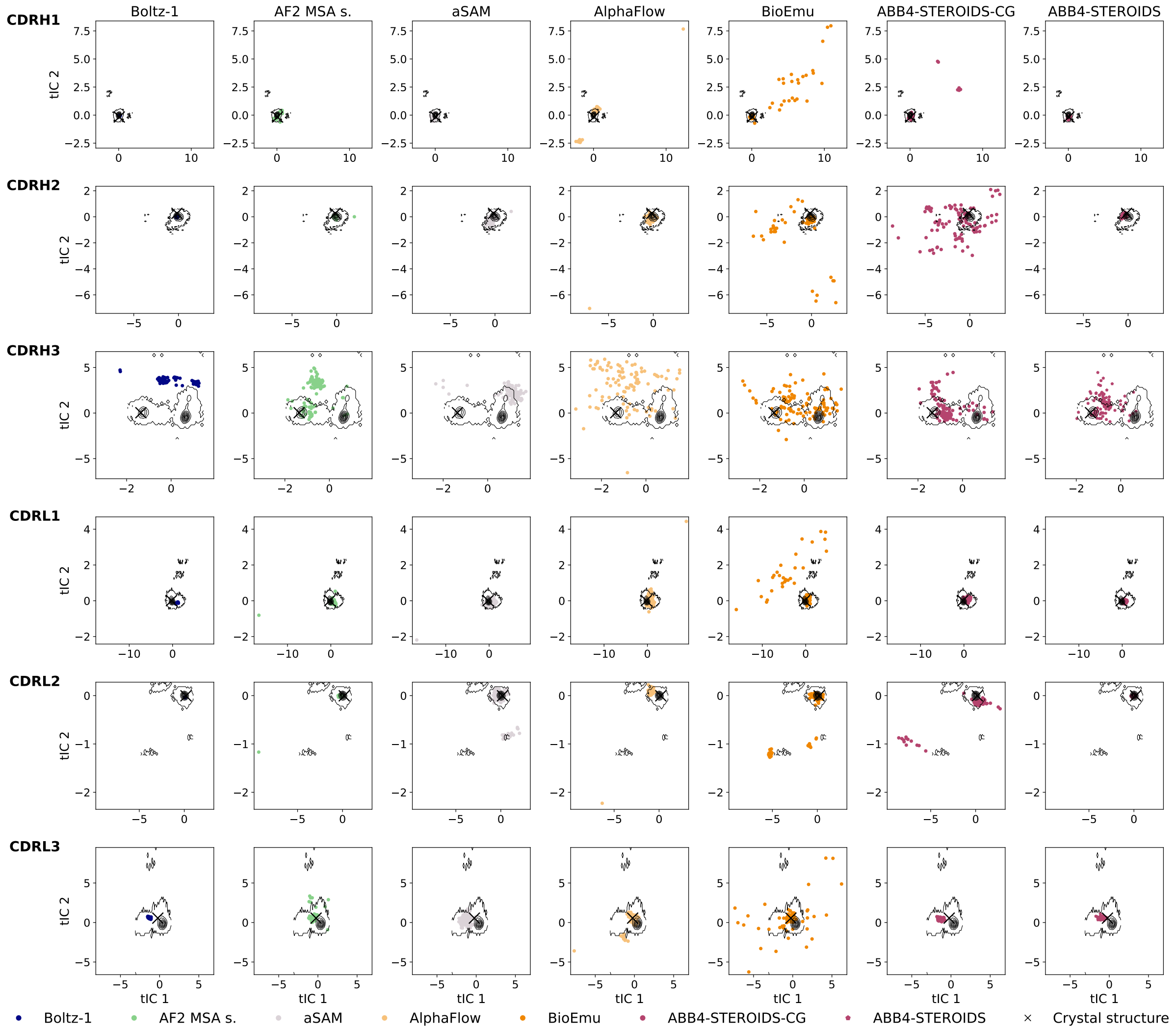

Figure S12: Comparison of the conformational ensemble sampled for test set antibody PDB 8oni. The first two time-lagged independent components (tICs) are plotted. The black contour indicates the all atom MD distributions. Predicted structures were mapped to the tICs and are represented by the coloured dots. The crystal structure from which the simulations was started is shown by the black cross

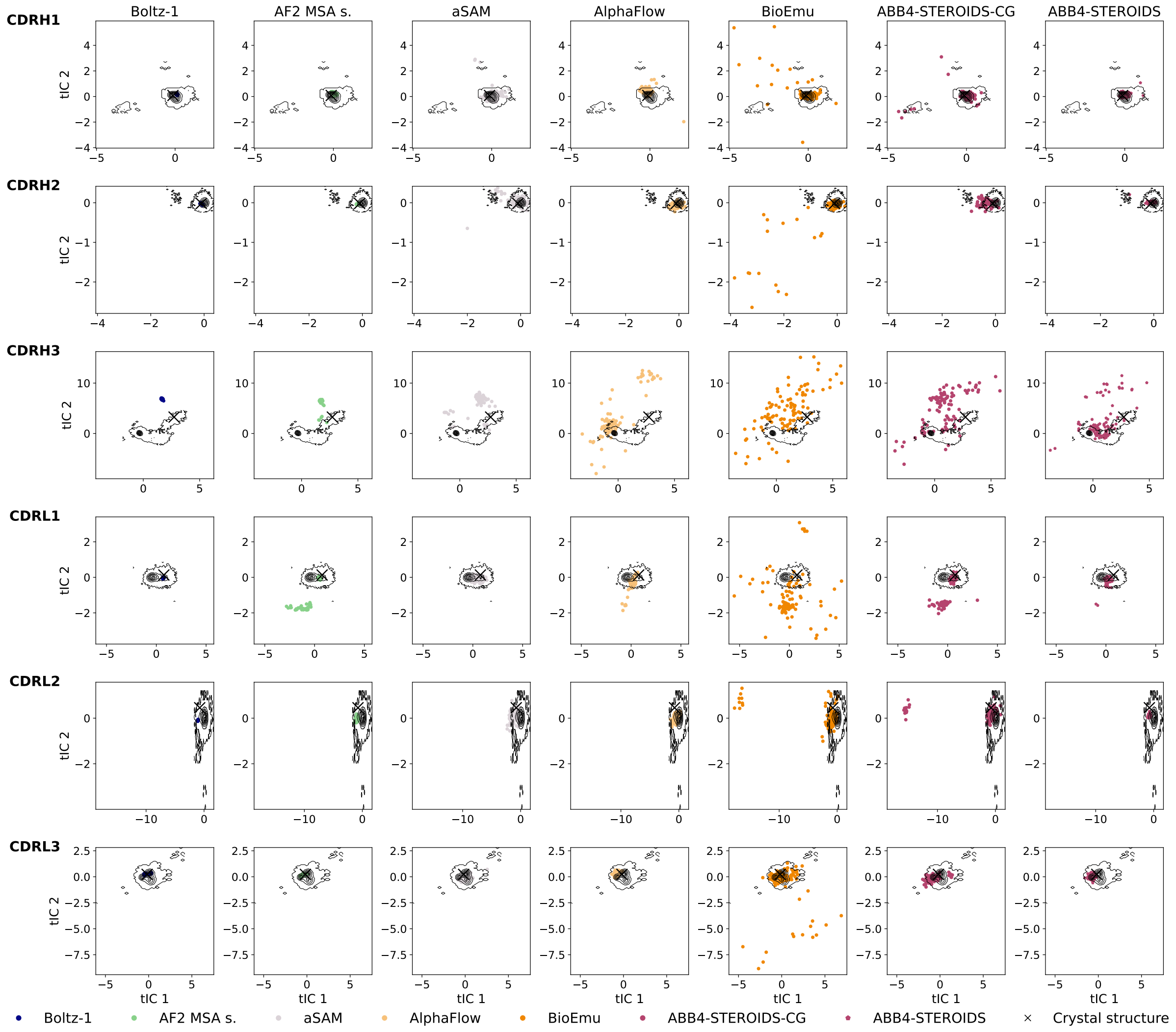

Figure S13: Comparison of the conformational ensemble sampled for test set antibody PDB 8r1d. The first two time-lagged independent components (tICs) are plotted. The black contour indicates the all atom MD distributions. Predicted structures were mapped to the tICs and are represented by the coloured dots. The crystal structure from which the simulations was started is shown by the black cross

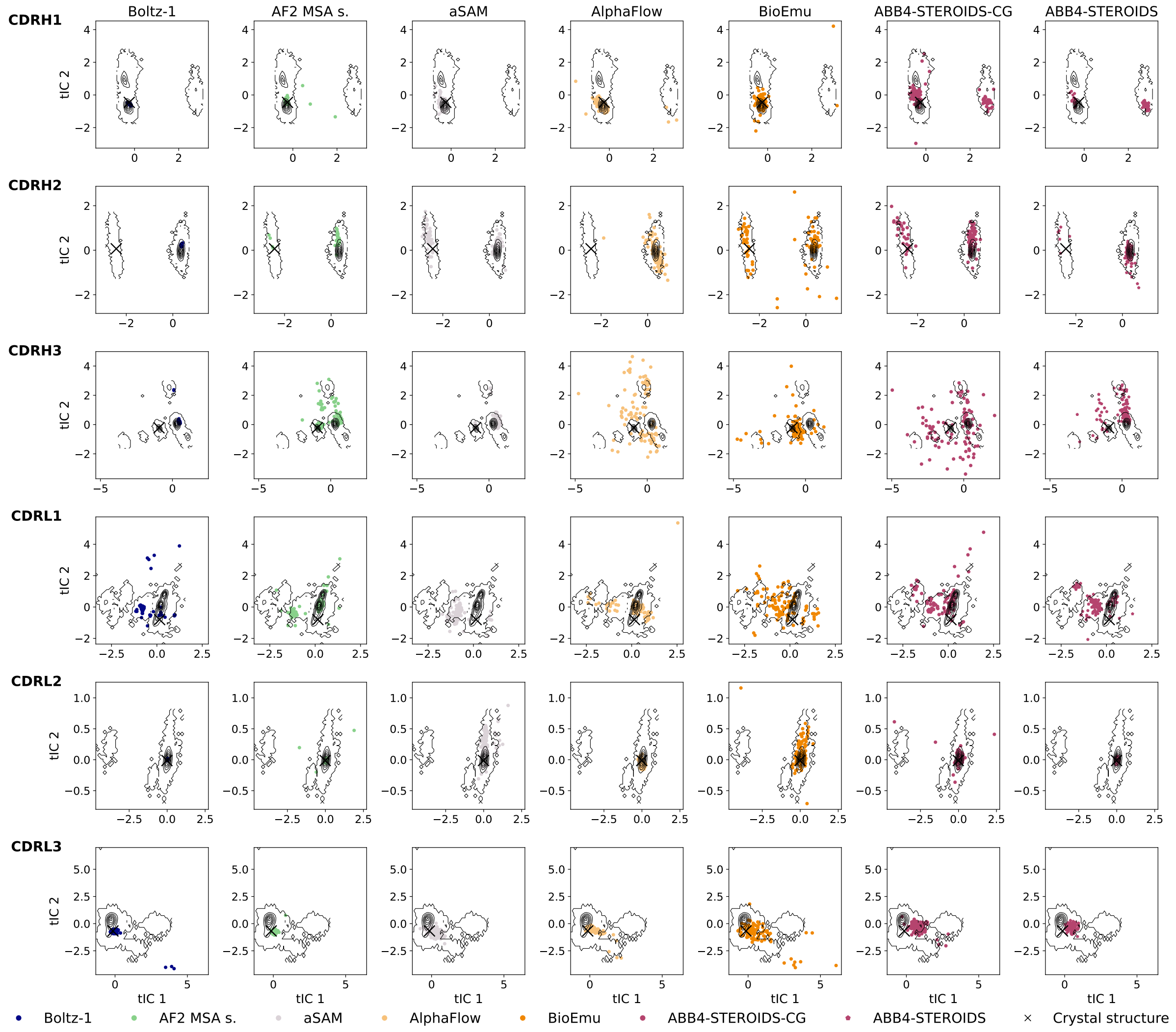

Figure S14: Comparison of the conformational ensemble sampled for test set antibody PDB 8vvb. The first two time-lagged independent components (tICs) are plotted. The black contour indicates the all atom MD distributions. Predicted structures were mapped to the tICs and are represented by the coloured dots. The crystal structure from which the simulations was started is shown by the black cross

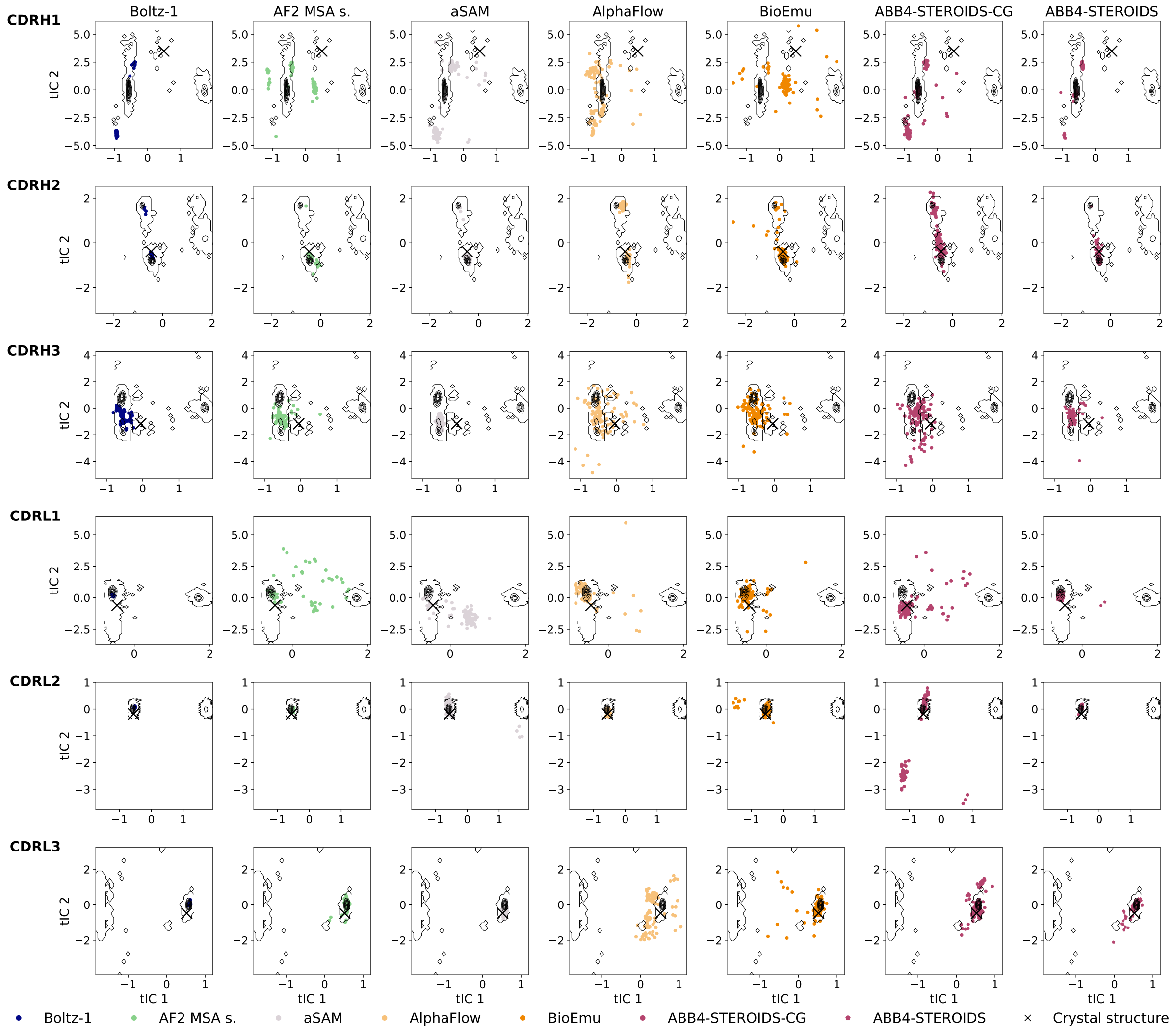

Figure S15: Comparison of the conformational ensemble sampled for test set antibody PDB 8w83. The first two time-lagged independent components (tICs) are plotted. The black contour indicates the all atom MD distributions. Predicted structures were mapped to the tICs and are represented by the coloured dots. The crystal structure from which the simulations was started is shown by the black cross

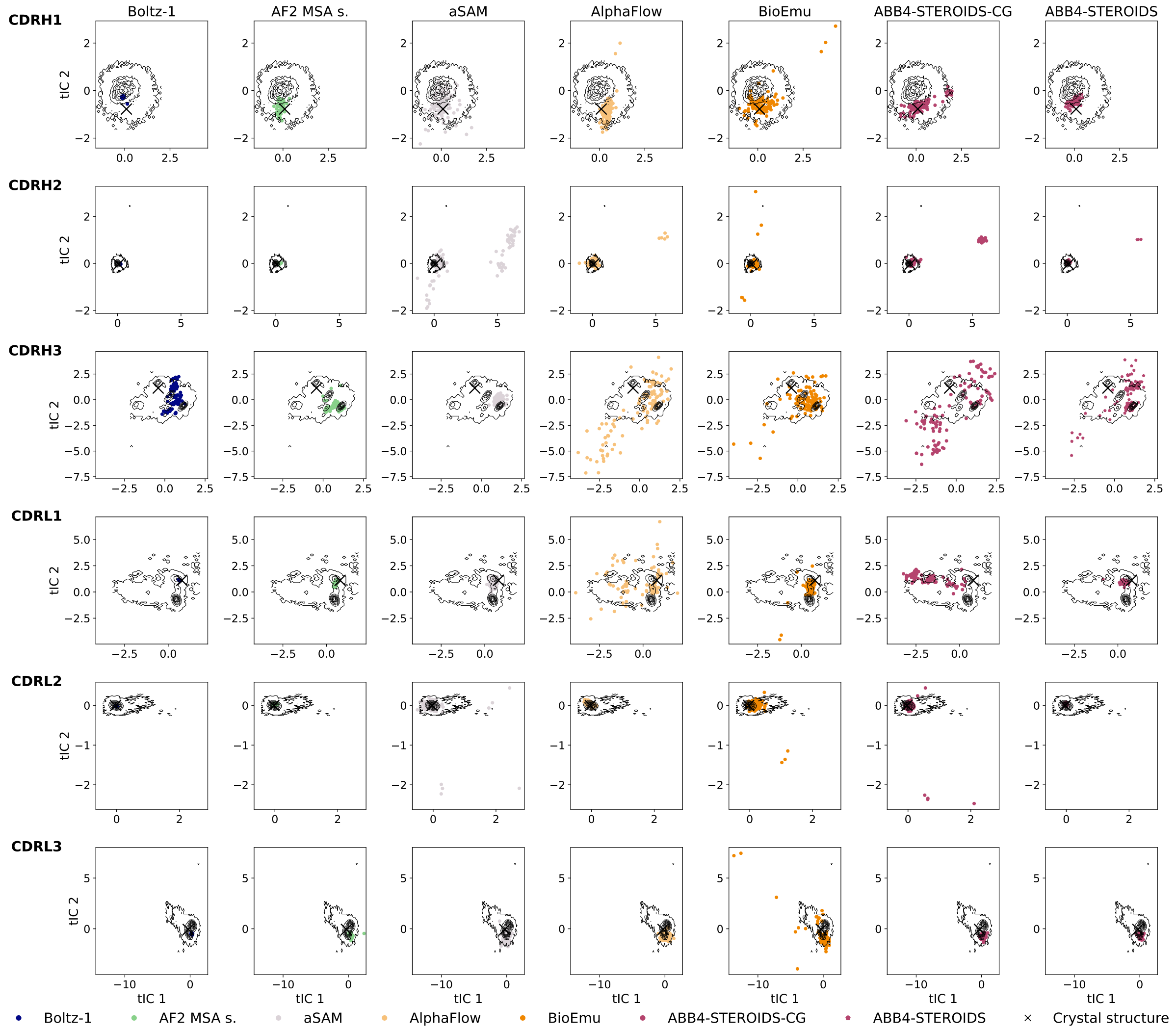

Figure S16: Comparison of the conformational ensemble sampled for test set antibody PDB 8gkj. The first two time-lagged independent components (tICs) are plotted. The black contour indicates the all atom MD distributions. Predicted structures were mapped to the tICs and are represented by the coloured dots. The crystal structure from which the simulations was started is shown by the black cross

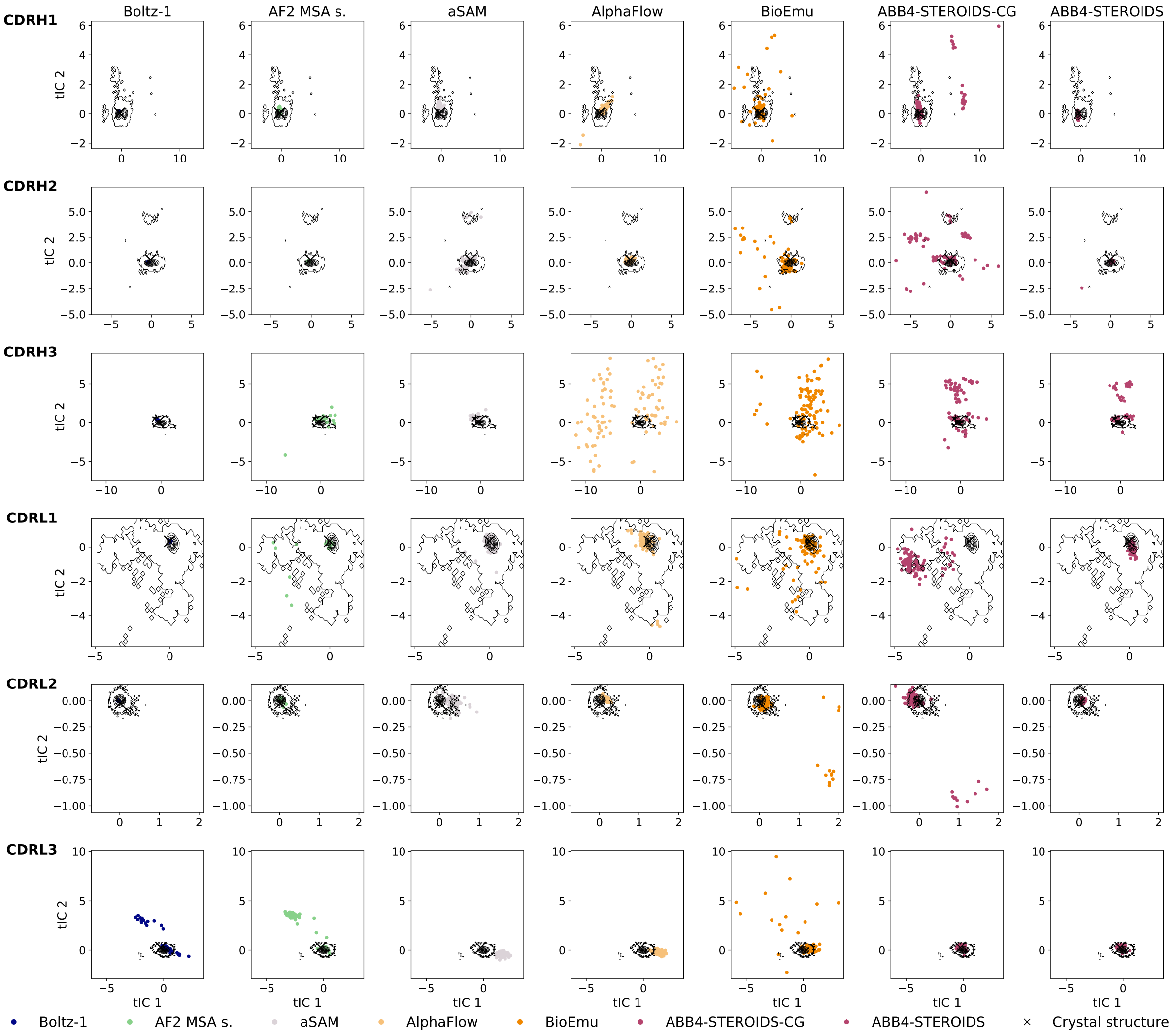

Figure S17: Comparison of the conformational ensemble sampled for test set antibody PDB 8jyr. The first two time-lagged independent components (tICs) are plotted. The black contour indicates the all atom MD distributions. Predicted structures were mapped to the tICs and are represented by the coloured dots. The crystal structure from which the simulations was started is shown by the black cross

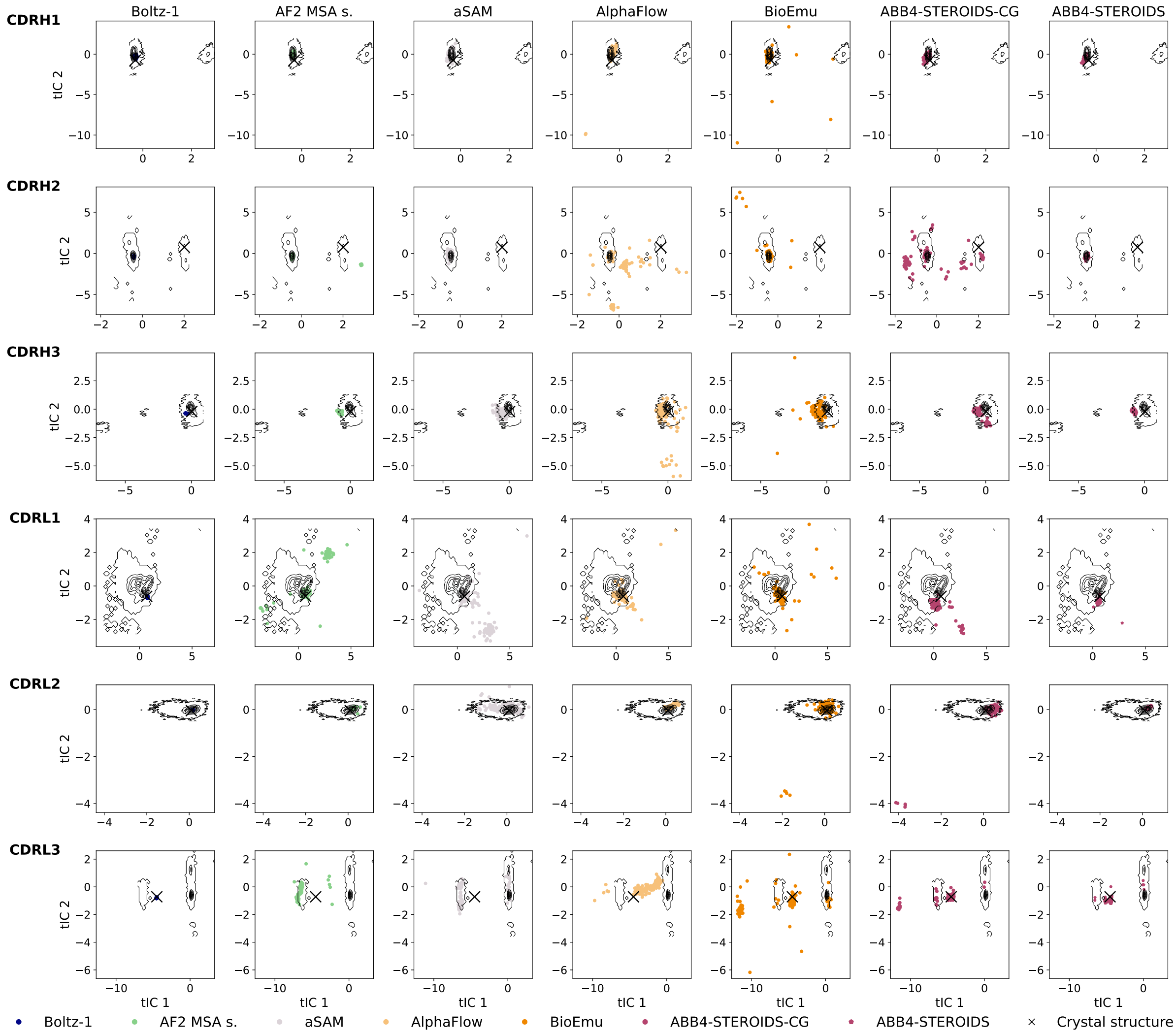

Figure S18: Comparison of the conformational ensemble sampled for test set antibody PDB 8vui. The first two time-lagged independent components (tICs) are plotted. The black contour indicates the all atom MD distributions. Predicted structures were mapped to the tICs and are represented by the coloured dots. The crystal structure from which the simulations was started is shown by the black cross

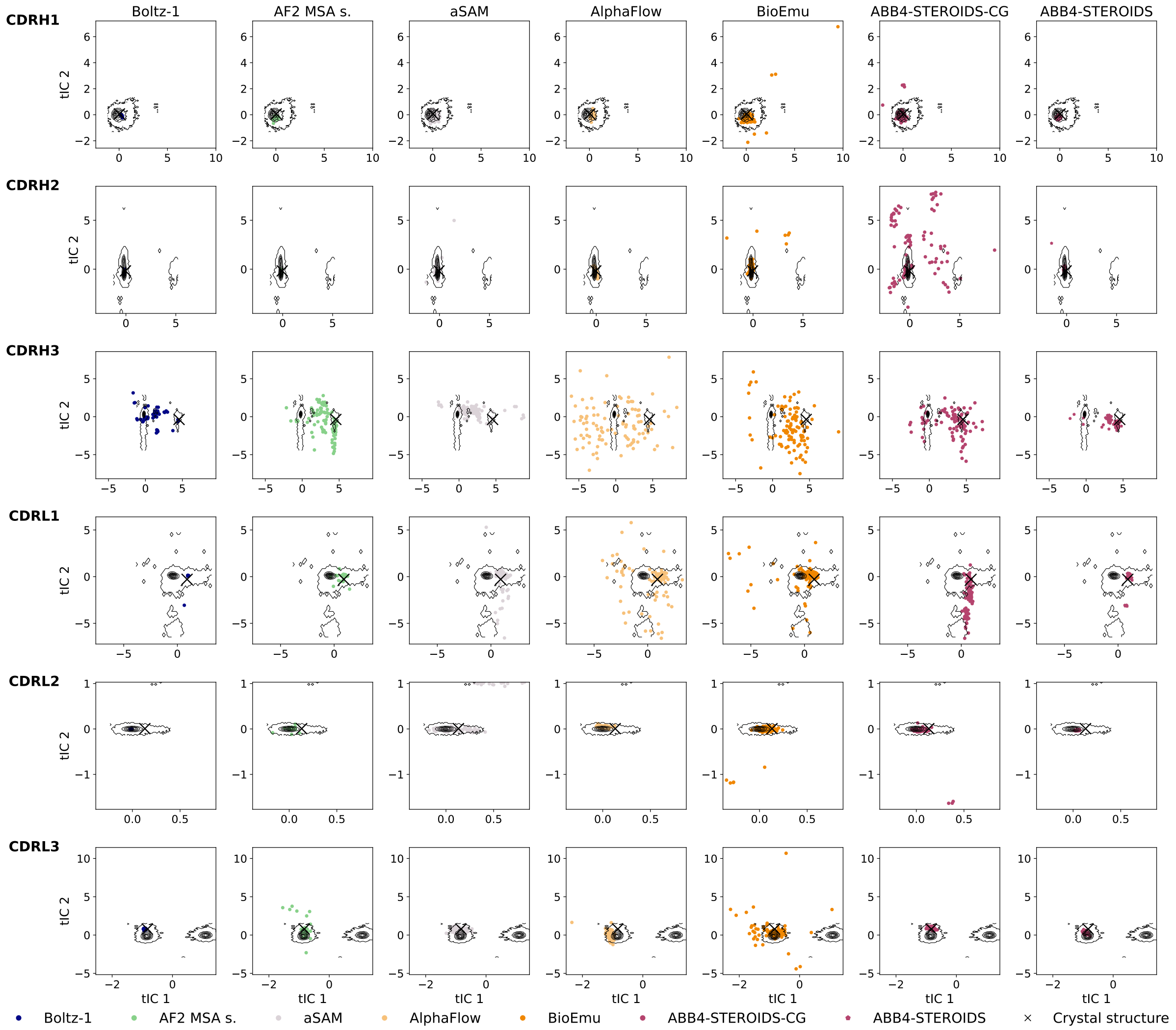

Figure S19: Comparison of the conformational ensemble sampled for test set antibody PDB 8w9h. The first two time-lagged independent components (tICs) are plotted. The black contour indicates the all atom MD distributions. Predicted structures were mapped to the tICs and are represented by the coloured dots. The crystal structure from which the simulations was started is shown by the black cross

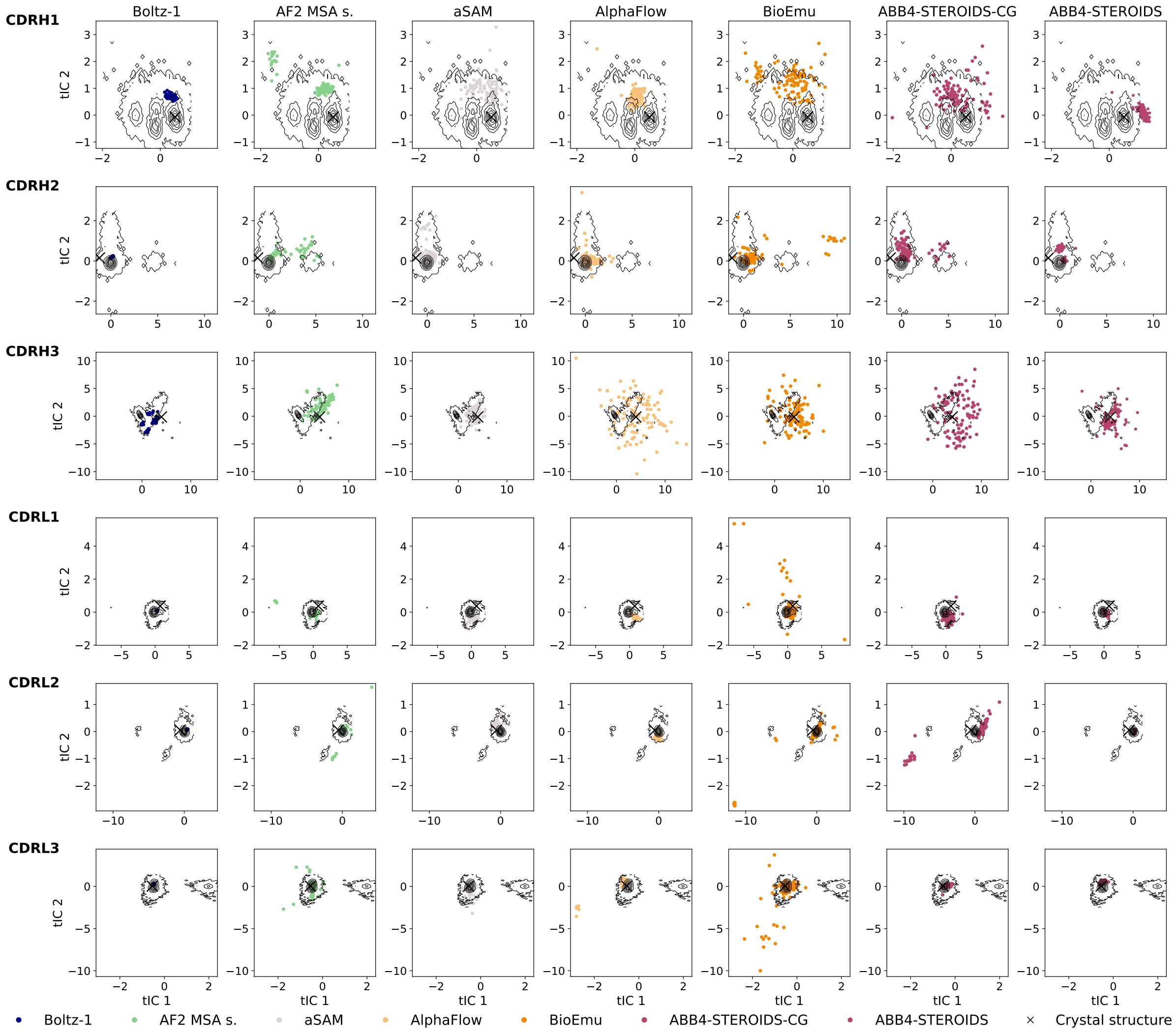

Figure S20: Comparison of the conformational ensemble sampled for test set antibody PDB 8xnh. The first two time-lagged independent components (tICs) are plotted. The black contour indicates the all atom MD distributions. Predicted structures were mapped to the tICs and are represented by the coloured dots. The crystal structure from which the simulations was started is shown by the black cross

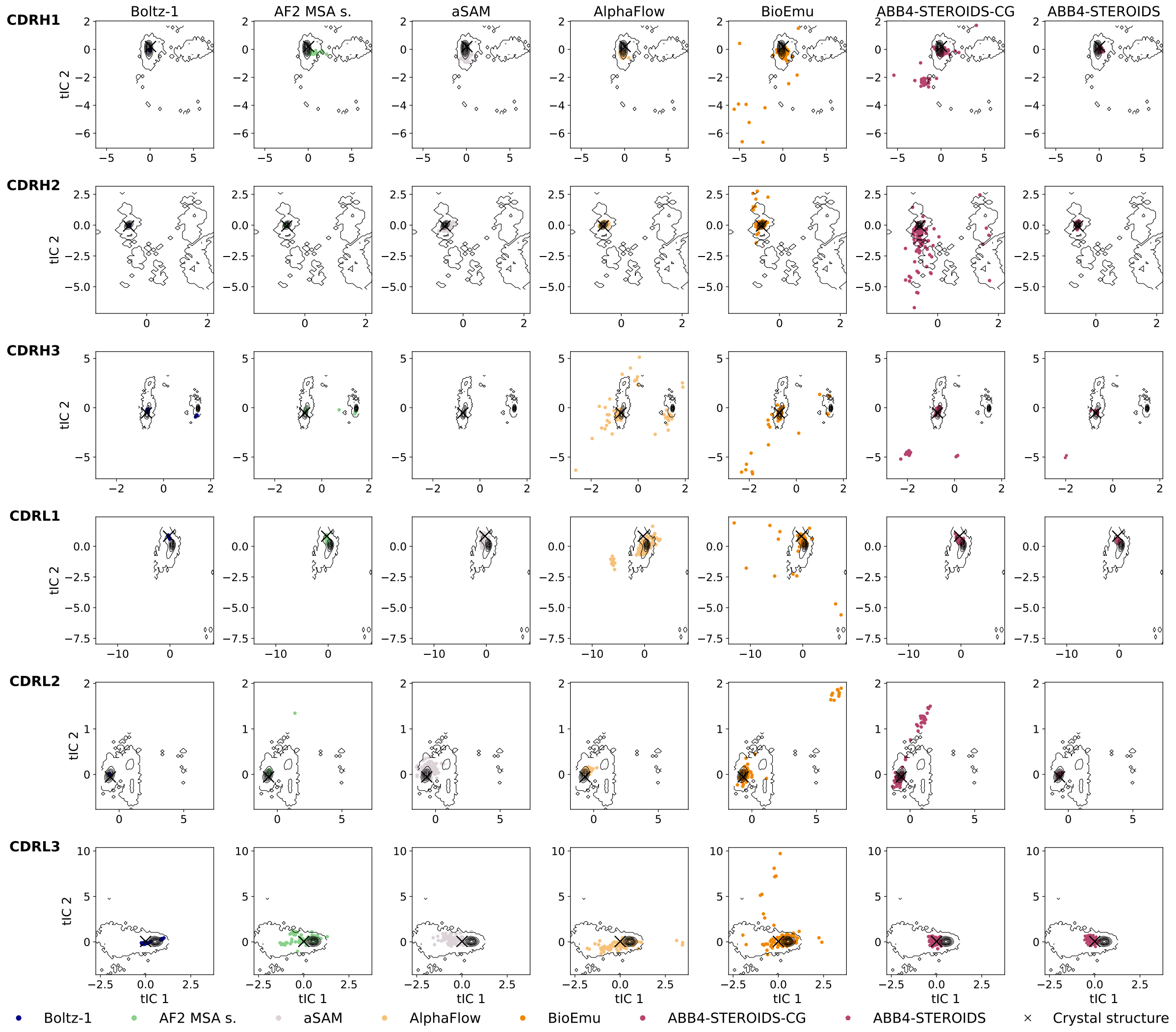

Figure S21: Comparison of the conformational ensemble sampled for test set antibody PDB 9g6s. The first two time-lagged independent components (tICs) are plotted. The black contour indicates the all atom MD distributions. Predicted structures were mapped to the tICs and are represented by the coloured dots. The crystal structure from which the simulations was started is shown by the black cross

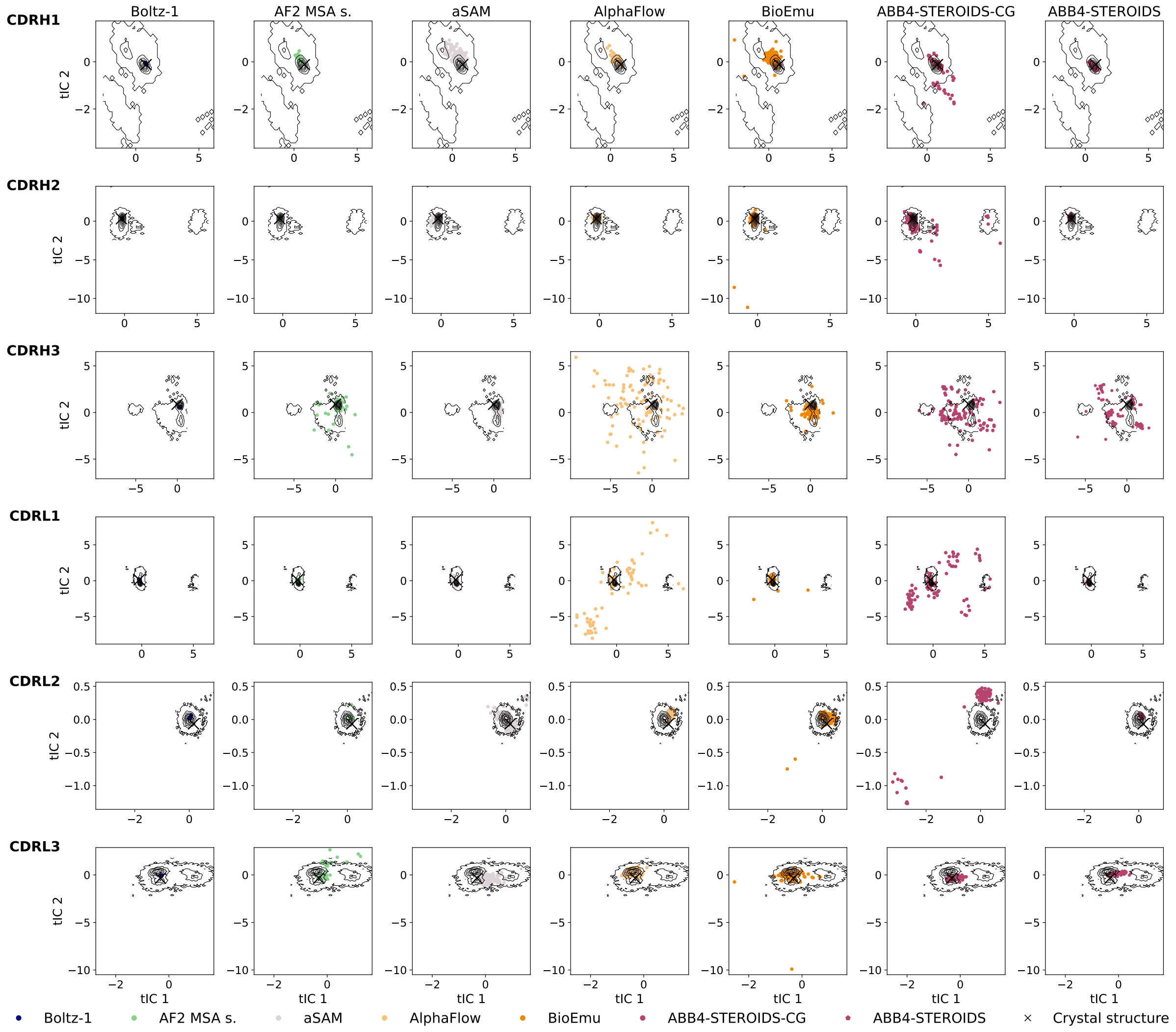

Figure S22: Comparison of the conformational ensemble sampled for test set antibody PDB 9h4r. The first two time-lagged independent components (tICs) are plotted. The black contour indicates the all atom MD distributions. Predicted structures were mapped to the tICs and are represented by the coloured dots. The crystal structure from which the simulations was started is shown by the black cross

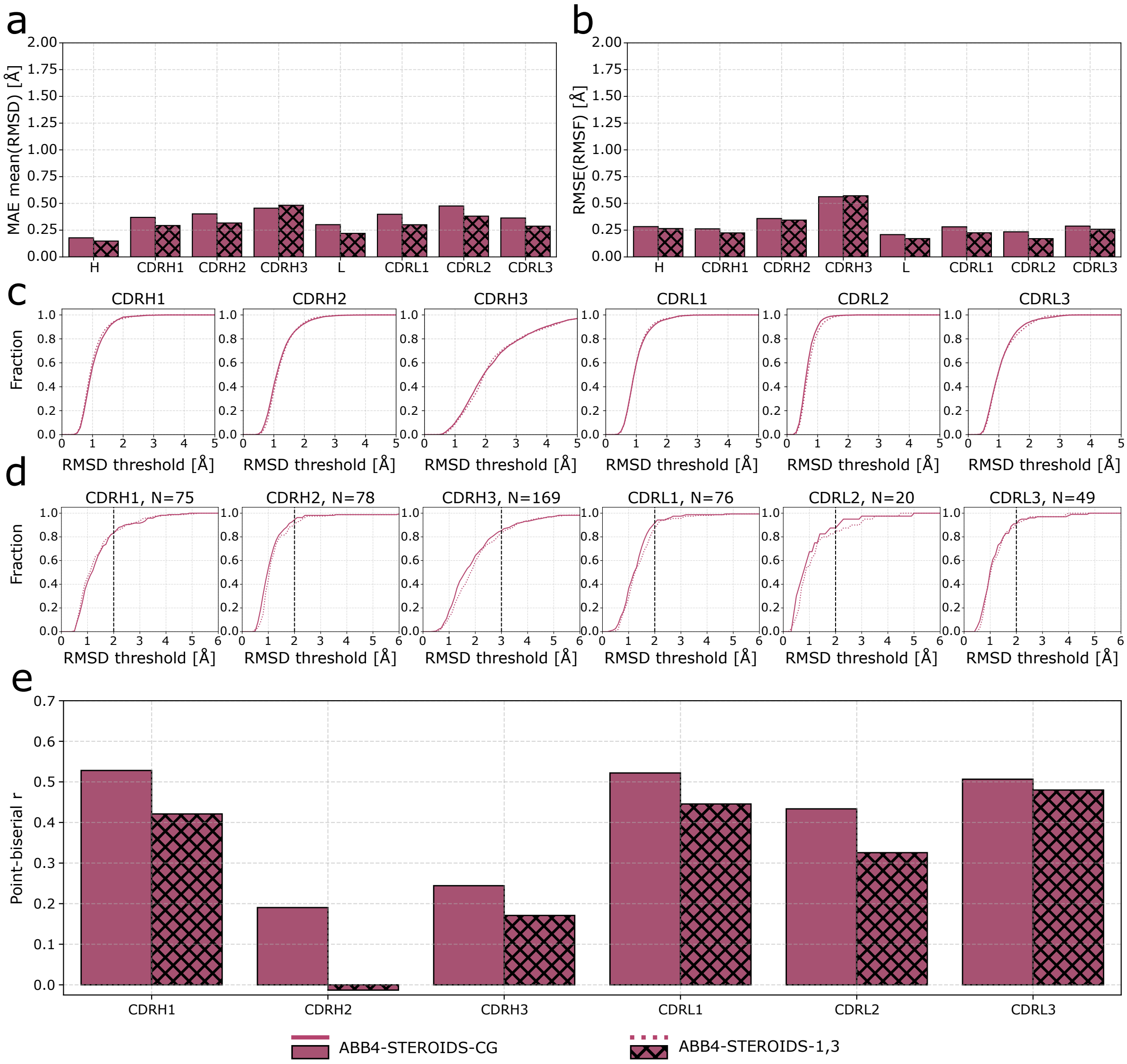

Figure S23: Evaluation of training stage ablation. In contrast to ABB4-STEROIDS-CG, trained with stages 1 to 3, ABB4-STERODIS-1,3 was only trained with stages 1 and 3. Mean absolute error (MAE) of RMSD (a) and root mean square error (RMSE) of RMSF (b) in the predicted ensembles compared to the MD simulations. Values computed across the heavy chain (H), light chain (L) and individual CDRs. c) Coverage of MD conformational frames as a function of distance threshold. d) Conformation coverage. Fraction of conformations observed in experimental ensembles covered as a function of distance. Dashed lines highlight ABB4-STEROIDS/-CG covering the highest number of conformations at given distance thresholds. e) Flexibility correlation. Point-biserial correlation coefficient (r) between binary flexibility labels assigned to experimental ensemble and RMSD in predicted ensembles for all CDR loops.

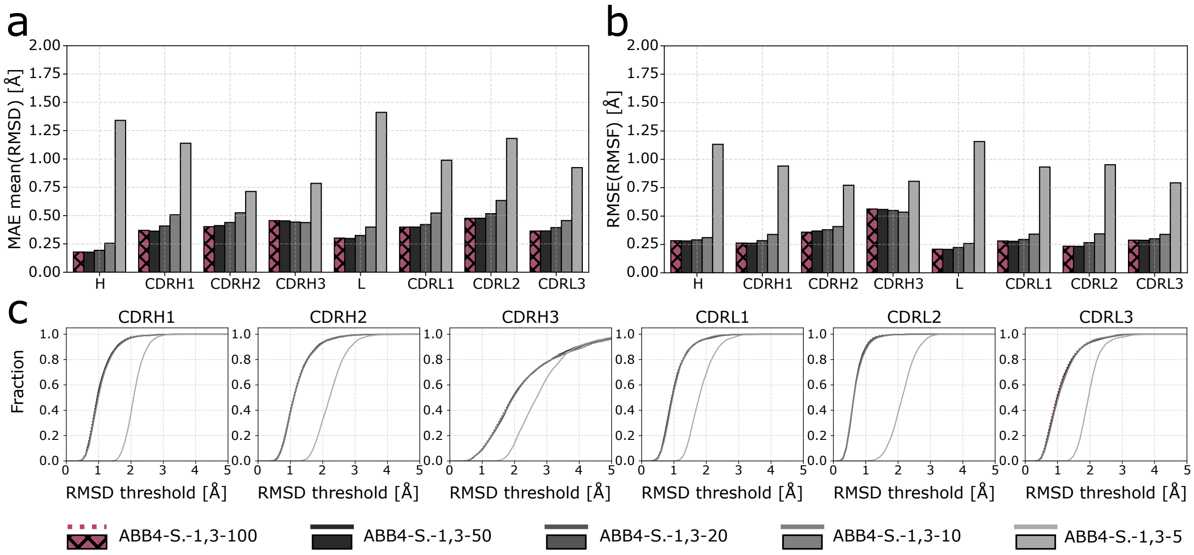

Figure S24: Evaluation of inference steps on model performance. ABB4-STEROIDS-1,3 with default inference procedure of 100 steps (ABB4-S.-1,3-100) and a reduced number of 50 (ABB4-S.-1,3-50), 20 (ABB4-S.-1,3-20), 10 (ABB4-S.-1,3-10) and 5 steps (ABB4-S.-1,3-5). Mean absolute error (MAE) of RMSD (a) and root mean square error (RMSE) of RMSF (b) in the predicted ensembles compared to the MD simulations. Values computed across the heavy chain (H), light chain (L) and individual CDRs. c) Coverage of MD conformational frames as a function of distance threshold.

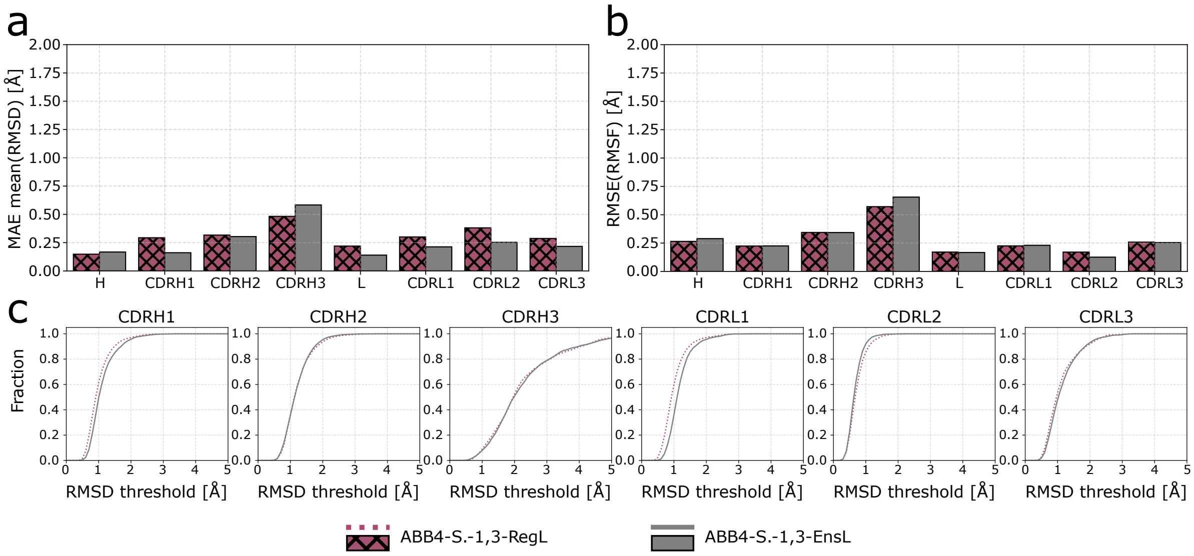

Figure S25: Evaluation of ensemble loss function on model performance. ABB4-STEROIDS-1,3 trained with the default loss (ABB4-S.-1,3-RegL) and ensemble loss (ABB4-S.-1,3-EnsL). Mean absolute error (MAE) of RMSD (a) and root mean square error (RMSE) of RMSF (b) in the predicted ensembles compared to the MD simulations. Values computed across the heavy chain (H), light chain (L) and individual CDRs. c) Coverage of MD conformational frames as a function of distance threshold.
